# Approximate Bayesian computation of transcriptional pausing mechanisms

**DOI:** 10.1101/748210

**Authors:** Jordan Douglas, Richard Kingston, Alexei J. Drummond

## Abstract

At a transcriptional pause site, RNA polymerase (RNAP) takes significantly longer than average to transcribe the nucleotide before moving on to the next position. At the single-molecule level this process is stochastic, while at the ensemble level it plays a variety of important roles in biological systems. The pause signal is complex and invokes interplay between a range of mechanisms. Among these factors are: non-canonical transcription events – such as backtracking and hypertranslocation; the catalytically inactive intermediate state hypothesised to act as a precursor to backtracking; the energetic configuration of basepairing within the DNA/RNA hybrid and of those flanking the transcription bubble; and the structure of the nascent mRNA. There are a variety of plausible models and hypotheses but it is unclear which explanations are better.

We performed a systematic comparison of 128 kinetic models of transcription using approximate Bayesian computation. Under this Bayesian framework, models and their parameters were assessed by their ability to predict the locations of pause sites in the *E*. *coli* genome.

These results suggest that the structural parameters governing the transcription bubble, and the dynamics of the transcription bubble during translocation, play significant roles in pausing. This is consistent with a model where the relative Gibbs energies between the pre and posttranslocated positions, and the rate of translocation between the two, is the primary factor behind invoking transcriptional pausing. Whereas, hypertranslocation, backtracking, and the intermediate state are not required to predict the locations of transcriptional pause sites. Finally, we compared the predictive power of these kinetic models to that of a non-explanatory statistical model. The latter approach has significantly greater predictive power (AUC = 0.89 cf. 0.73), suggesting that, while current models of transcription contain a moderate degree of predictive power, a much greater quantitative understanding of transcriptional pausing is required to rival that of a sequence motif.

**Author summary:** Transcription involves the copying of a DNA template into messenger RNA (mRNA). This reaction is implemented by RNA polymerase (RNAP) successively incorporating nucleotides onto the mRNA. At a transcriptional pause site, RNAP takes significantly longer than average to incorporate the nucleotide. A model which can not only predict the locations of pause sites in a DNA template, but also explain *how* or *why* they are pause sites, is sought after.

Transcriptional pausing emerges from cooperation between several mechanisms. These mechanisms include non-canonical RNAP reactions; and the thermodynamic properties of DNA and mRNA. There are many hypotheses and kinetic models of transcription but it is unclear which hypotheses and models are required to predict and explain transcriptional pausing.

We have developed a rigorous statistical framework for inferring model parameters and comparing hypotheses. By applying this framework to published pause-site data, we compared 128 kinetic models of transcription with the aim of finding the best models for predicting the locations of pause sites. This analysis offered insights into mechanisms of transcriptional pausing. However, the predictive power of these models lacks compared with non-explanatory statistical models - suggesting the data contains more information than can be satisfied by current quantitative understandings of transcriptional pausing.

## Introduction

The reaction pathways (Fig 1) of transcription elongation have been studied extensively. RNA polymerase (RNAP) exists inside a transcription bubble that translocates along the double stranded DNA. Within the polymerase, the messenger RNA (mRNA) and template DNA form a DNA/RNA hybrid [1–7]. Throughout the main transcription elongation pathway, RNAP successively alternates between the **pretranslocated** and **posttranslocated** positions, employing a Brownian ratchet mechanism [8, 9]. In order to bind and incorporate the next nucleotide onto the 3′ end of the mRNA, the active site must be accessible and RNAP must be in the posttranslocated position.

**Fig 1.**
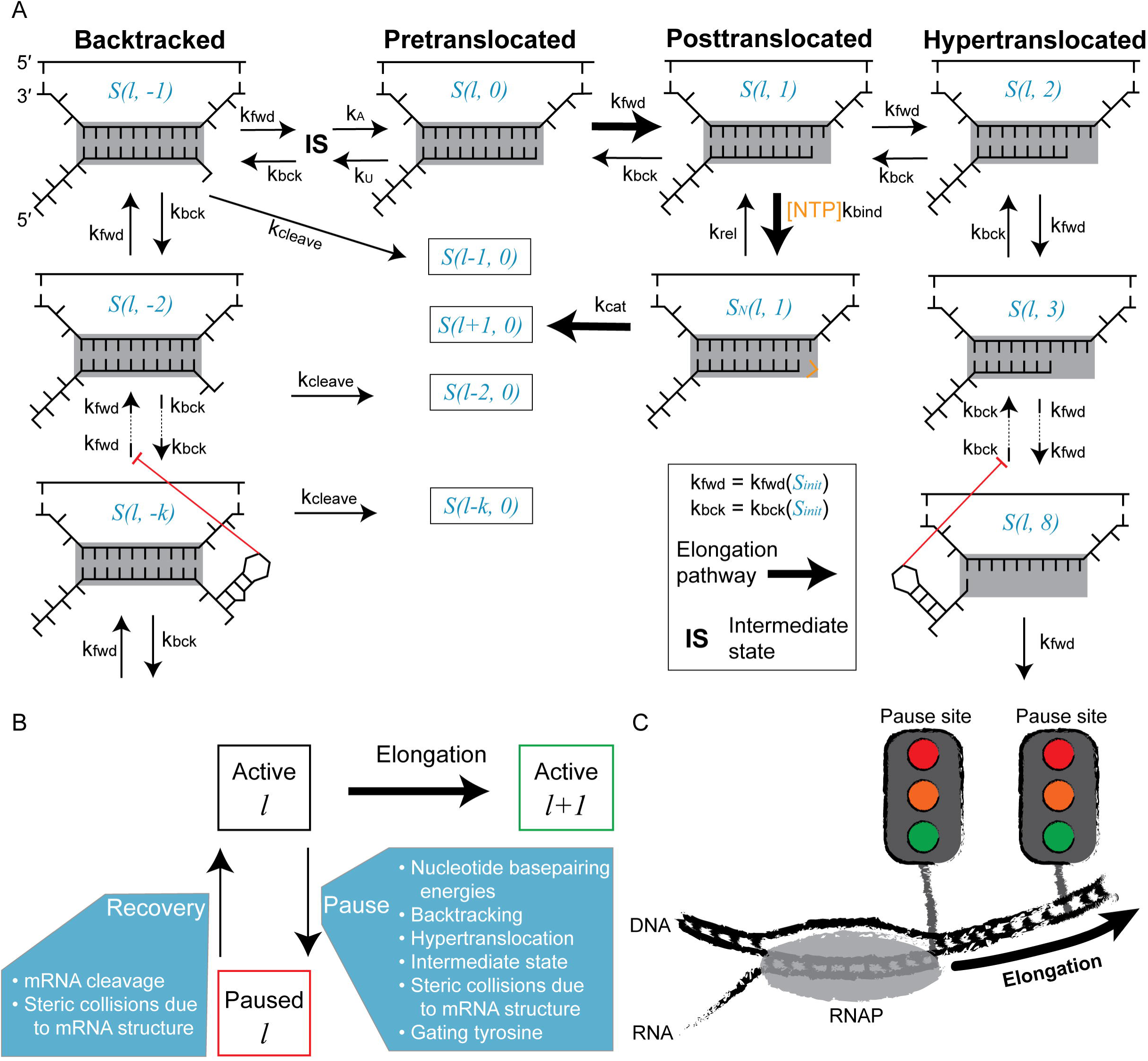
The kinetics of transcriptional pausing. A: State diagram of transcription elongation (bold arrows) and its off-pathway events. RNA polymerase (grey rectangle) translocates the template in the 3^′^ → 5^′^ direction. State notation described in main text. Inhibitory effects that RNA secondary structures have on translocation are displayed. B: Schematic depiction of pausing, including mechanisms and effectors of pausing and pause recovery described in A. C: Pause sites are analogous to traffic lights. If the light is green, the RNA polymerase may go. If the light is red it must wait an indefinite amount of time for the light to turn green. The colour of the upcoming traffic light and the time until it turns green are random variables.

However, RNAP does not always conform to this catalytically productive pathway. As off-pathway events, RNAP can **backtrack** or **hypertranslocate**. When backtracking (Fig 1A, LHS), RNAP translocates upstream (in the 3′ direction along the template) [9–11]. This causes the 3′ end of the mRNA to exit the RNA polymerase through the NTP entry channel thus rendering the polymerase active site inaccessible. Recovery to the elongation pathway is stimulated by transcription elongation factors, such as GreA in prokaryotes and S-II in eukaryotes, which cleave the 3′ mRNA [11–13]. It can also occur through RNAP intrinsic cleavage activity [13]. When hypertranslocating (Fig 1A, RHS), RNAP moves 5′-bound down the template such that the active site is beyond the 3′ mRNA [14, 15]. Each nucleotide by which RNAP hypertranslocates causes the shortening of the DNA/RNA hybrid by one basepair. Hypertranslocation is a key pathway to intrinsic termination [16], whereby an upstream mRNA hairpin destabilises RNAP and induces termination [17].

It has been hypothesised that a catalytically inactive **intermediate state** (IS) acts as a precursor for backtracking [1, 18–22]. Under this hypothesis, this state, sometimes also known as the inactive state [23] or the elemental paused elongation complex [22], has taken many forms in the literature and exists as an intermediate between the pretranslocated and backtracked states (Fig 1). Entry into this state from a pretranslocated complex is achieved by rearrangement of the RNAP trigger-loop and fraying of the 3′ mRNA, in a manner that inhibits elongation but not translocation [24].

For prokaryotes and eukaryotes, transcription elongation rates range from 20-120 bp/s [25–27]. However individual RNAP molecules proceed quite erratically along their template. Approximately once every 100 bp [23, 28] there exists a pause site which takes significantly longer to transcribe, oftentimes on the timescale of seconds or minutes [19, 29, 30]. Extended pauses can lead to RNA polymerase traffic jams [31, 32]. For the most part, pause sites are likely to be detrimental to the organism.

Nonetheless, transcriptional pausing plays a range of important biological roles in certain systems [33].

### 1. Gene expression

For example, the 5′ UTR of HIV-1 contains a pause site immediately downstream from the TAR hairpin [19, 34]. TAR is a regulatory element that upregulates transcription upon binding to viral protein Tat [35]. It is therefore beneficial for the virus if there exists a pause site downstream from TAR, as this gives Tat a greater temporal opportunity to bind to TAR before RNAP has left the proximity.

### 2. Modulating RNA folding

For example, the *ribDEAHT* operon of *Bacillus subtilis* encodes genes involved in riboflavin synthesis [36]. The *ribD* riboswitch, which manifests in the nascent mRNA, can adopt either the terminator fold (eliciting termination) or the antiterminator fold (enabling transcription). The former is favoured in the presence of flavin mononucleotide. Transcriptional pause sites flank the riboswitch, thus providing this ligand a greater opportunity to apply its effect [37].

### 3. RNA splicing

RNA splicing involves the pairing of a donor and an acceptor splice site. As this process occurs cotranscriptionally [38], the chance of the cellular splicing machinery selecting any given donor-acceptor pair is dependent on transcription elongation velocity and transcriptional pausing [39–41]. The positioning and strength of pause sites therefore contributes to the proportions of splice variants in systems which employ alternative splicing.

The dwell time at a pause site is approximately exponentially distributed [19, 23, 30] and subsequently any given pause site can be quantified by how likely the RNAP is to pause, and an escape time half-life for when it does pause [19, 23]. While comparisons can be made between pause sites and stop signs or speed bumps, due to their stochastic nature pause sites are more akin to traffic lights (Fig 1C).

However the mechanisms that elicit pausing to occur in the first place are complex and multipartite [19, 22, 24]. A range of sequence-dependent factors contribute to the pause signal. Among the known effectors: 1) when the DNA/RNA hybrid of a posttranslocated state is weak relative to that of the pretranslocated state, RNAP may favour the pre state over the post state thus leading to a pause [21]. This delay can facilitate backtracking, which further extends the pause [42]. 2) The DNA up to 24 bp downstream from a pause site can have an effect on pausing [43]. This is unexpected because the DNA in this region is too far from RNAP to be disrupted by translocation. This phenomenon may be caused by the downstream DNA bending in a sequence-dependent manner [19, 43]. 3) It is expected that the secondary structure of the mRNA inhibits translocation [1, 44]. Upstream structures could inhibit backwards translocation and downstream structures could inhibit forward translocation. 4) Pauses occur more frequently when there is a pyrimidine at the 3′ end of the mRNA [28, 45]. This could be due to nucleotide-specific interactions between the mRNA and the protein [22, 45]. 5) The four NTPs have different dissociation constants *K*_*D*_ and different rates of catalysis *k*_*cat*_ [46], which can give the four nucleotides different pause propensities. 6) Nucleotide misincorporations destabilise the enzyme and can elicit pausing [47–49].

A consensus sequence of the pause site for the *E*. *coli* RNAP has recently been identified [28, 50]. This motif reveals that the nucleotides at the 3′ and 5′ ends of the hybrid are important, as well as the incoming NTP which is usually a GTP. By comparing a nucleotide sequence to a motif [50], one may be able to predict the locations of pause sites. However it still leaves much unsaid about the physical processes that govern pausing. Although sequence-dependent explanatory models have been applied [1, 45, 46, 51], a systematic and large scale analysis of the accuracy of this approach has not yet been done.

It would be greatly beneficial to have a model for both predicting the locations of and explaining the mechanisms behind pause sites, for any arbitrary gene sequence. There are still uncertainties pertaining to the mechanism behind transcriptional pausing we would like to resolve. 1) To what extent does mRNA secondary structure inhibit the translocation thereby modifying the pausing behaviour? Does using a prediction of the mRNA structure enhance the model [1, 44]? 2) Does utilising knowledge of the gated tyrosine residue that inhibits translocation between the backtrack-1 and backtrack-2 states improve the model [11]? It could be the case that the backtrack-1 state is readily accessible and incorporating this feature into the model improves its accuracy. 3) Some [1, 18–22], but not all [44, 51, 52], models have been built with the inclusion of the IS that RNAP must enter before backtracking. Is this model feature essential to explain the sequence-specific properties of pausing or is it a redundant feature that introduces unnecessary complexity into the model?

By virtue of the availability of data from a high-throughput detection of pause sites across the entire *E*. *coli* transcriptome by Larson et al. 2014 [28], we were able to explore these model variants. In this study we used a Bayesian approach to interrogate this dataset. The volume of this dataset allowed us to 1) evaluate how reliable this modelling approach can be for the prediction of pause sites, and 2) select the best model, and its parameters, to better understand the mechanics of transcriptional pausing.

## Models

We explored two approaches for predicting pause sites: 1) the simulation of kinetic models as continuous-time Markov processes (based on the kinetic scheme shown in Fig 1), and 2) by using a simple naive Bayes classifier. Both models were trained on the aforementioned dataset [28].

The first approach involves stochastic simulation of transcription at the single-molecule level using the Gillespie algorithm [53, 54]. This is done in a similar fashion to our previous work [55], but here we have used the model to predict the dwell time at each site instead of mean velocity under force.

### Preliminaries

Let *S2*(*l, t*) denote a state, where *l* is the current length of the mRNA and *t* ∈ℤ is the position of the polymerase active site with respect to the 3′ end of the mRNA (Fig 1A). *t* = 0 when pretranslocated and *t* = 1 when posttranslocated. *t* < 0 denotes backtracked states and *t >* 1 denotes hypertranslocated states.

Standard Gibbs free energies Δ_*r*_*G*^0^(= Δ*G*) involved in duplex formation are used to calculate forward and backward translocation rates. These terms are approximated using nearest neighbour models. The total Gibbs energy of state *S* – arising from nucleotide basepairing and dangling ends – is

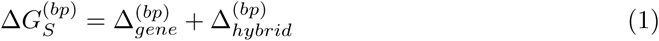

where SantaLucia’s DNA/DNA basepairing parameters [56] are used to calculate 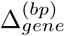 and Sugimoto’s DNA/RNA parameters [57] are used for 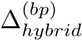. For the latter, dangling end energies are estimated as described by Bai et al. 2004 [51] and only apply to the posttranslocated position. Δ*G* terms are expressed relative to the thermal energy of the system, in units of *k*_*B*_*T*, where *k*_*B*_*T* = 0.6156 kcal/mol at *T* = 310 K.

### Comparing kinetic models

To better understand the mechanisms that govern transcriptional pausing, we not only estimated the kinetic model parameters but also the kinetic model itself. In this section we describe six different model settings. Each model setting has a discrete set of values giving a cross-product of 2 × 2 × 4 × 2 × 2 × 2 = 128 models to compare.

#### Inclusion of the intermediate state, 𝕀𝕊

There has been discussion concerning whether there exists an IS that acts as an entry point for backtracking from the elongation pathway [1, 22]. The IS is catalytically inactive and can act as a prolonged pause state regardless of whether backtracking is subsequently instigated [42]. While incorporating this physical process may offer additional explanatory power to the model, two additional parameters *k*_*U*_ and *k*_*A*_ must be estimated. We therefore compared two models: a general model (𝕀𝕊 = 1) where the IS exists and is necessary for backtracking, and a simpler model (𝕀𝕊 = 0) where there is no IS and RNAP can backtrack freely (and there are two fewer parameters to estimate).

#### Inclusion of the gating tyrosine, 𝔾𝕋

The crystal structure of the *S*. *cerevisiae* Pol II complex by Cheung et al. 2011 [11] reveals a gating tyrosine that may inhibit backtracking. While back translocation into the *S*(*l*, −1) position may be permitted, further backtracking into *S*(*l*, −2) is likely delimited by this amino acid. We were interested whether the gating tyrosine plays is an effector of transcriptional pausing, so we compared two models: one model (𝔾𝕋 = 1) where RNAP can readily translocate between *S*(*l*, 0) and *S*(*l*, −1) but translocation between *S*(*l*, − 1) and *S*(*l*, −2) is much less favourable, and a simpler model (𝔾𝕋 = 0) where the effects of the gating tyrosine are ignored (same as Fig 1).

#### Estimating the translocation transition state, 𝕋𝕊

As the transcription bubble migrates along the gene, so too do the basepairing configurations within the DNA/RNA hybrid and the DNA/DNA gene. In order for RNAP to translocate from one position into the next, disruption of one hybrid basepair and one gene basepair may be required. This translocation is assumed to occur through a translocation transition state [55]. Four sequence-dependent methods for estimating this transition state are described (Fig 2A).

**Fig 2.**
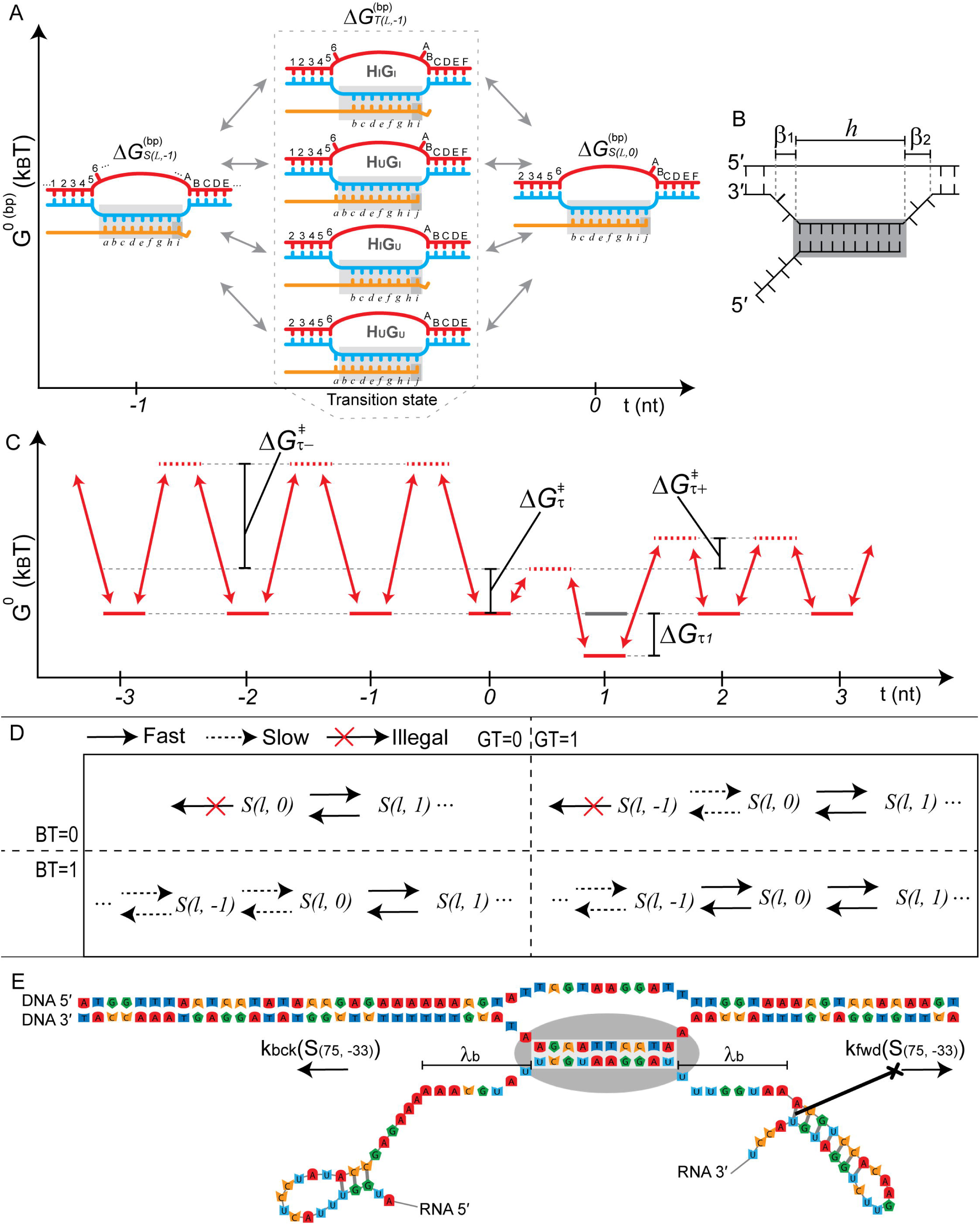
Kinetic model variants and parameters. A: The four translocation transition state models. Four possible transition states between the backtracked *S*(*l*, −1) and pretranslocated *S*(*l*, 0) states are displayed. A transition state can comprise of hybrid (gene) basepairs that are the union or the intersection of the hybrid: *H*_*U*_ and *H*_*I*_ respectively (gene: *G*_*U*_ and *G*_*I*_ respectively). B: The transcription bubble is described by three parameters. In this example *β*_1_ = 2, *h* = 10, *β*_2_ = 3. C: Gibbs energy landscape of translocation with the energies of translocation states *S* (solid lines) and translocation transition states *T* (dashed lines) shown. The displayed energies are sequence-independent in this diagram: the energies from nucleic acid thermodynamic parameters would be added onto these values in the final calculations. Figure corresponds to model 𝕀𝕊 = 0, 𝔾𝕋 = 0, 𝔹𝕋 = 1, ℍ𝕋 = 1. D: State diagrams showing the relationship between 𝔾𝕋 and 𝔹𝕋, where 𝕀𝕊 = 0. Dashed line arrows refer to translocation steps that are augmented by 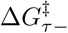. E: RNA secondary structures can act as blockades (ℝ𝔹 = 1) and prohibit translocation. The effect of *λ*_*b*_ = 8 nt is shown for the first 75 nucleotides of the *rpoB* gene.

The DNA/RNA basepairs within the hybrid of the translocation transition state are assumed to be a subset of the basepairs within the two neighbouring states. These basepairs could be the union ∪ or the intersection ∩of the two sets. Similarly, the DNA/DNA basepairs within the gene are assumed to be either the union ∪ or the intersection ∪ of the DNA/DNA basepairs within the two neighbouring states.

This gives a total of four models. For ease of pronunciation, union is denoted by a subscripted *U* and intersection is denoted by a subscripted *I*. In the model 𝕋𝕊 = *H*_*I*_*G*_*I*_, the basepairs within the transition state’s hybrid (*H*) and gene (*G*) are each the intersection of the two respective neighbouring states. Following the same notation, the other three models are 𝕋𝕊 = *H*_*I*_*G*_*U*_, 𝕋𝕊 = *H*_*U*_ *G*_*I*_, and 𝕋𝕊 = *H*_*U*_ *G*_*U*_. These four models describe different mechanisms of translocation and could give different sequence-dependent emergent properties.

To determine which, if any, of these four models are the most suitable we estimated the value of 𝕋𝕊.

#### Inclusion of RNA secondary structure blockades, ℝ𝔹

The simplest models of incorporating RNA secondary structure as a translocation inhibitor make the assumption that transcription is sufficiently slow for mRNA to fold into its minimum free energy structure within the same timescale [1, 44]. As a more complex model, one could invoke a kinetic model of cotranscriptional folding derived from something to the likes of Kinfold [58]. It was of interest how much this first model, and its questionable assumptions, contributed to predictive and explanatory power with respect to transcriptional pausing. We therefore compared two models of RNA folding: the model ℝ𝔹 = 0 with no RNA folding and the model ℝ𝔹 = 1 where the minimum free energy structure, as predicted by ViennaRNA suite [59, 60], is used as a translocation blockade.

#### Modelling off-pathway translocation, 𝔹𝕋 + ℍ𝕋

It is unclear to what extent backtracking 𝔹𝕋 ∈ {0, 1} and hypertranslocation ℍ𝕋 ∈ {0, 1} play roles in transcriptional pausing. While these pathways are certainly real observed phenomena, it may be the case that they are not required to adequately explain pausing. To elucidate this, we explored models in which either backtracking, or hypertranslocation, or both, or neither, are illegal pathways.

### Parameterisation of the kinetic model

In this section we describe how the rates presented in Fig 1 are calculated. The parameters for calculating these rates were in some cases held constant based off previous estimates, while in other cases were estimated from the data. Where estimated, a prior distribution is required. Table 1 summarises these constants and priors.

**Table 1.**
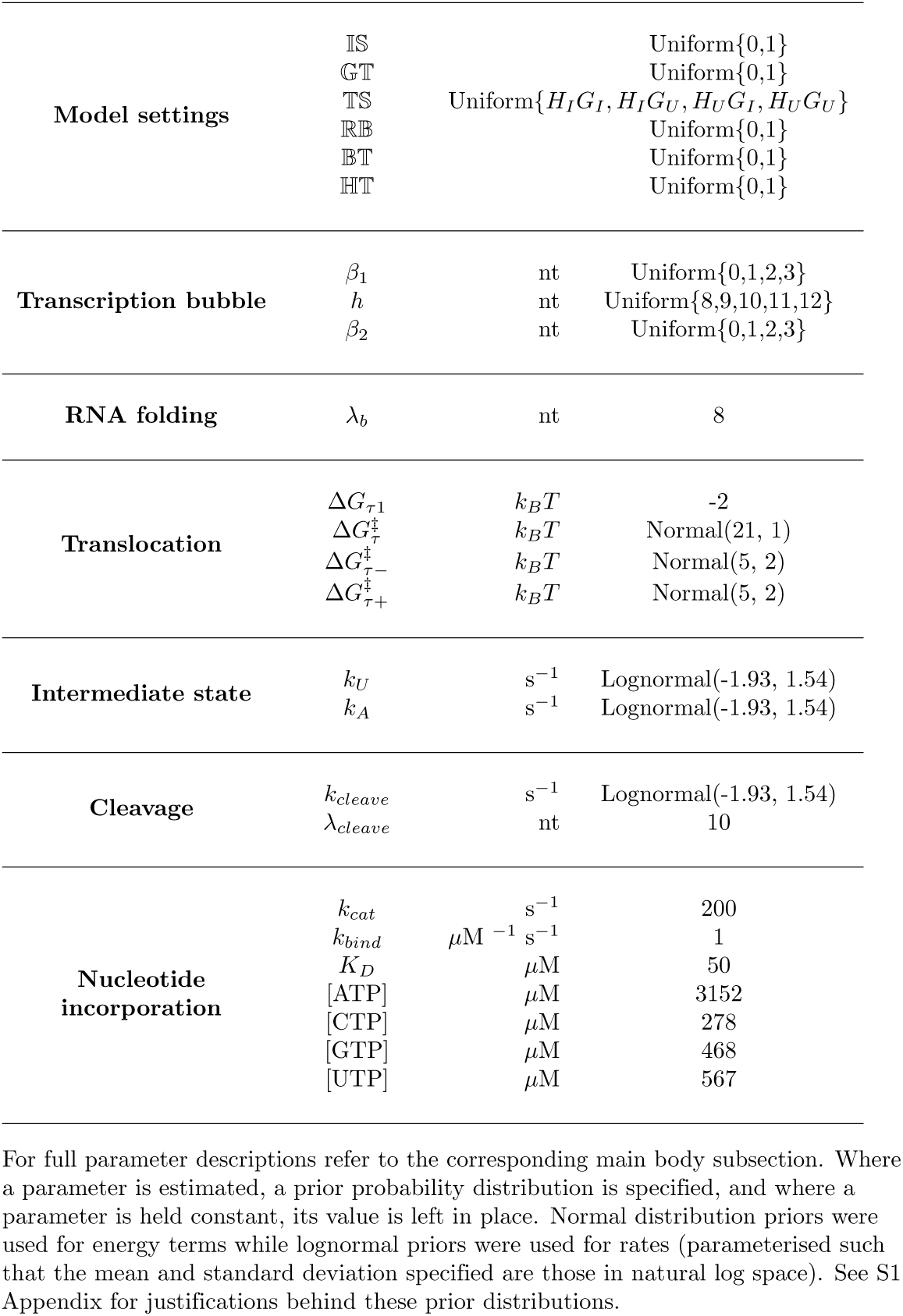
Summary of parameters used in the kinetic model.

#### Transcription bubble

The transcription bubble is described by three parameters (Fig 2B): the number of unpaired template nucleotides on the 3′ and 5′ ends of the bubble, *β*_1_ and *β*_2_ respectively, and the number of paired template nucleotides in the DNA/mRNA hybrid, *h* [1]. These three parameters are to be estimated from the data and expected to have a profound effect on the sequence-dependent properties of translocation [51].

#### RNA folding

RNA folding is incorporated into the translocation model in a similar fashion to Tadigotla et al. 2006 [44] where, provided that ℝ𝔹 = 1, the RNA minimum free energy (MFE) structure is predicted using the ViennaRNA suite [59, 60], and these structures can block translocation. However, the *λ*_*b*_ nucleotides proximal to the mRNA entry and exit channels respectively are assumed to be unable to adopt a secondary structure due to steric collisions with the enzyme, and are therefore unable to block translocation.

Suppose that (*r*_1_, *r*_2_, … *r*_*l*_) is the mRNA sequence of state *S*(*l, t*). Let *R*(*i, j*), where 1 ≤ *i* ≤ *j* ≤ *l* + 1, be the subsequence of nucleotides (*r*_*i*_, *…, r*_*j-*1_) which are basepaired in the MFE structure of mRNA subsequence. *R*(*i, i*) is the empty set. Let *U* (*l, t*) and *D*(*l, t*) be the set of basepaired nucleotides in the mRNA, upstream and downstream of RNAP respectively.

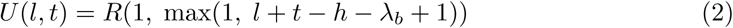

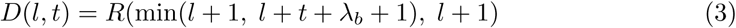

Then, the backwards translocation rate out of state *S*(*l, t*) is equal to zero, due to steric collision with upstream RNA, if 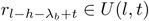. Similarly, the forward translocation rate is equal to zero, due to downstream collisions, if 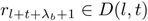.

#### Translocation

The computation of translocation rates invokes transition state theory and is parameterised as an extension to our previous work [55]. Translocation from state *S* is described by the rate of backward translocation *k*_*bck*_(*S*) and the rate of forward translocation *k*_*fwd*_(*S*). *k*_*bck*_ and *k*_*fwd*_ are therefore functions of *S* and are sequence-dependent. As the transcription bubble migrates along the gene, the basepairing within the gene and within the hybrid changes with it.

Translocation rates are principally computed from thermodynamic properties of nucleic acid duplexes [56, 57]. In order for RNA polymerase to translocate forward (backward), two basepairs must be disrupted: (1) the basepair at the downstream (upstream) edge of the transcription bubble, and (2) the basepair at the upstream (downstream) end of the DNA/mRNA hybrid. The strength of basepairing in these regions affects the sequence-dependent rate of translocation. For example in SantaLucia’s parameters [56] 5′ TA/AT is the weakest doublet at − 0.94 *k*_*B*_*T* while 5′ GC/CG is the strongest at Δ*G*^(*bp*)^ = −3.64 *k*_*B*_*T*. A larger energy barrier must be overcome to disrupt the latter.

Let *T* (*l, t*) be the translocation transition state between states *S*(*l, t*) and *S*(*l, t* + 1). Using transition state theory, the rates of forward and backwards translocation out of *S*(*l, t*) are

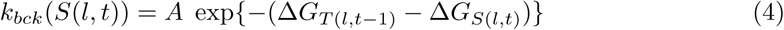

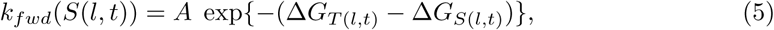

for some pre-exponential constant *A*, arbitrarily set here to 10^6^ s^*-*1^ [55]. Two energetic terms are required to compute these rates; the Gibbs energy of the translocation state Δ*G*_*S*(*l,t*)_, and the Gibbs energy of the translocation transition state Δ*G*_*T* (*l,t*)_. We will describe these two components separately.

First, calculating Δ*G*_*S*(*l,t*)_ is straightforward (Fig 2C). It requires one translocation parameter – Δ*G*_*τ*1_. Δ*G*_*S*(*l,t*)_ is primarily computed from DNA/DNA and DNA/mRNA Gibbs energies 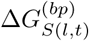 of the state.

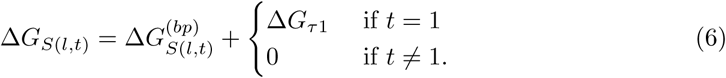

where Δ*G*_*τ*1_ is a term added onto the Gibbs energy of basepairing of the posttranslocated state. Δ*G*_*τ*1_ was found to be a necessary parameter to describe elongation sufficiently for the *E*. *coli* RNAP and was estimated as ∼−2 *k*_*B*_*T* [55] and is set accordingly throughout this study (Table 1).

Second, calculating Δ*G*_*T* (*l,t*)_ is more complex (Fig 2C). It requires a method for estimating the nucleic acid energies of the transition state, and three translocation parameters – 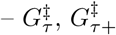, and 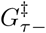.

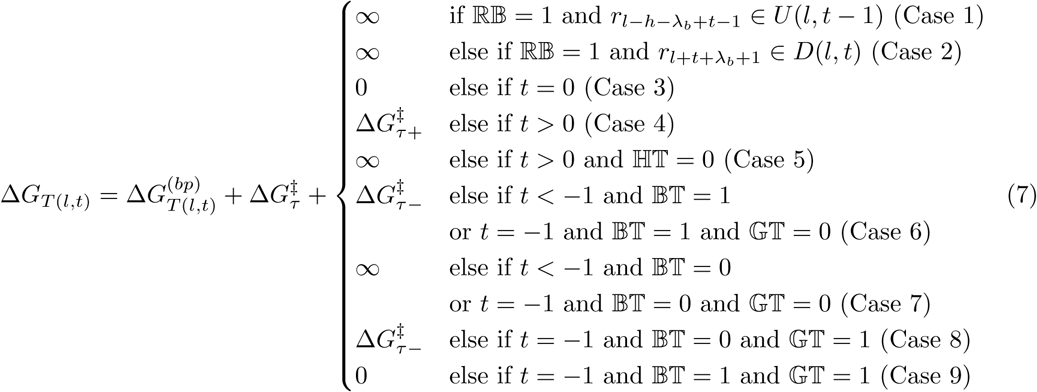

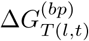 is computed entirely from nucleic acid parameters [56, 57] and is dependent on the value of 𝕋𝕊. See **Translocation transition state**.

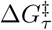 describes the intrinsic activation barrier of translocation that must be overcome in all translocation reactions. It should be set to a magnitude such that forward and backward translocation rates are in the order of 10^1^-10^2^ s^*-*1^ [1, 55, 61].

The remaining terms in Equation 7 are dependent on the position of RNAP (backtracked, pretranslocated, etc.) and the model itself and are broken down into cases.

Case 1: *Upstream RNA blockade*. When the RNA blockade model is enabled, ie. ℝ𝔹 = 1, backward translocation is not permitted if it requires breaking a basepair in the mRNA immediately upstream of the enzyme. In this case, the transition state of such a reaction has a Gibbs energy of infinity, corresponding to a translocation rate of 0. See **RNA folding**.

Case 2: *Downstream RNA blockade*. Analogous to Case 1. Forward translocation is not permitted if it requires breaking a downstream basepair.

Case 3: *Main elongation pathway*. *T* (0, *l*) is the transition state between the pre and posttranslocated states. The total energy of the transition state is 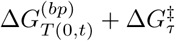 and acts as a baseline for all other activation energies.

Case 4: *Hypertranslocation*. When hypertranslocation is permitted, ie. ℍ𝕋 = 1, translocation into or out of a state where *t >* 0 is further augmented by the hypertranslocation energy barrier 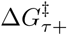.

Case 5: *Illegal hypertranslocation*. When hypertranslocation is disabled, ie. ℍ𝕋 = 0, then activation energies which correspond to hypertranslocation are set to infinity, corresponding to translocation rates of zero.

Case 6: *Backtracking*. When backtracking is enabled ie. 𝔹𝕋 = 1, then the backtracking activation energy 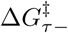 is added onto transition states associated with entering or leaving a backtracked state *S*(*l, t*) where *t* < −1. Furthermore, if the gating tyrosine is omitted from the model, ie. 𝔾𝕋 = 0, then this additional energy barrier also applies to the transition between *S*(*l*, 0) and *S*(*l, -*1). See Fig 2D for state diagrams of Cases 6-9.

Case 7: *Illegal backtracking*. When backtracking is disabled 𝔹𝕋 = 0, then RNAP may not translocate upstream of *S*(*l*, −1) so the energy of the transition state is infinity, and the translocation rate is zero. If the gating tyrosine is also omitted from the model 𝔾𝕋 = 0, then *S*(*l, -*1) is also an illegal state.

Case 8: *Slow backstepping*. When the gating tyrosine is modelled, but backtracking is omitted {𝔾𝕋 = 1, 𝔹𝕋 = 0}, translocation between *S*(*l*, 0) and *S*(*l*, −1) is permitted however is assumed to be a slow reaction and therefore the transition state energy is further augmented by 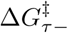.

Case 9: *Fast backstepping*. When the gating tyrosine and backtracking are both modelled {𝔾𝕋 = 1, 𝔹𝕋 = 1}, translocation between *S*(*l*, 0) and *S*(*l*, −1) is permitted and is assumed to be a fast reaction no different to that between *S*(*l*, 0) and *S*(*l*, 1). Backtracking beyond this point is still assumed to be slow (Case 6).

In summary, the rates of translocation are calculated using transition state theory (Equations 4 and 5). These rates are are sequence-dependent and are calculated from translocation state energies (Equation 6) and translocation transition state energies (Equation 7). These two terms are dependent on nucleic acid thermodynamic parameters [56, 57] and four model parameters 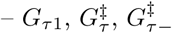, and 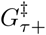. An estimate of the Gibbs energy of the translocation transition state 𝕋𝕊 is also required (Fig 2A), as well as the values of ℝ𝔹, ℍ𝕋, 𝔹𝕋, and 𝔾𝕋.

#### Translocation transition state

This section describes how 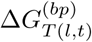 is computed, given the transition state model 𝕋𝕊.

Suppose that *S* is the current state. Let *b*_*i*_ be a basepair. Then, *b*_*i*_ is an element of *G*(*S*) if *b*_*i*_ is a DNA/DNA basepair in state *S*, and *b*_*i*_ is an element of *H*(*S*) if *b*_*i*_ is a DNA/RNA basepair in *S*.

Let *T* (*l, t*) be the translocation transition state between neighbouring states *S*(*l, t*) and *S*(*l, t* + 1). The set of basepairs in *H*(*T* (*l, t*)) and *G*(*T* (*l, t*)) depend on the current transition state model 𝕋𝕊.

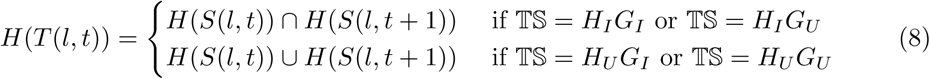

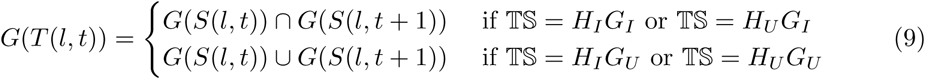

Once the set of basepairing nucleotides comprising the gene *G*(*T* (*l, t*)) and the hybrid *H*(*T* (*l, t*)) are known, the sequence-dependent Gibbs energy component of the translocation transition state 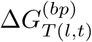 is readily computed [56, 57].

However, because the *number* of basepairs in each of the four transition state models 𝕋𝕊 differ, and their corresponding energies therefore systematically differ, 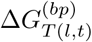 must be normalised such that the four models achieve the same average translocation rates.

This was achieved by calculating the mean value of 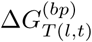 across all positions in the *E*. *coli rpoB* gene, for each transition state model 𝕋𝕊. The normalisation constants were calculated as −15.96 *k*_*B*_*T*, −11.26 *k*_*B*_*T*, −13.52 *k*_*B*_*T*, and −8.819 *k*_*B*_*T*, for models *H*_*I*_*G*_*I*_, *H*_*I*_*G*_*U*_, *H*_*U*_ *G*_*I*_, and *H*_*U*_ *G*_*U*_, respectively.

For example, in model 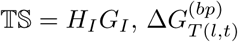 is equal to *-*15.96 *k*_*B*_*T* plus the actual basepairing energies of the transition state computed from nearest neighbour models [56, 57].

#### Intermediate state

Entry into the IS can be achieved by forward translocation from *S*(*l*, −1), or by inactivation from the pretranslocated state *S*(*l*, 0). This first pathway is a sequence-dependent reaction with a rate constant *k*_*fwd*_(*S*(*l, -*1)). This second pathway occurs at a constant rate described by parameter *k*_*U*_. Similarly the IS can be exited by backwards translocation into *S*(*l*, −1) with rate *k*_*bck*_(*S*(*l*, 0)), or by reactivation into *S*(*l*, 0) at a constant rate of *k*_*A*_. If 𝕀𝕊 = 0 the IS is omitted and translocation can directly bypass. See Fig 2D.

#### Cleavage

During cleavage, the 3′ end of the mRNA is, through some arbitrary mechanism, truncated. We used a similar model to Lisica et al. 2016 [13], where cleavage is described by two parameters: the first-order rate constant of cleavage *k*_*cleave*_ and the maximum number of positions RNAP can be backtracked by in order for cleavage to occur *λ*_*cleave*_.

Through cleavage, RNAP is restored from a backtracked state to a pretranslocated state.

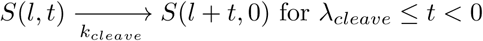

#### Nucleotide incorporation

Nucleotide incorporation is described by the rate of catalysis *k*_*cat*_, the second order rate constant of NTP binding *k*_*bind*_, and the NTP dissociation constant 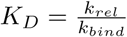, where *k*_*rel*_ is the rate of NTP release. Partial equilibrium approximations were not made in the NTP binding step.

### Worked example

In this section, the outbound rates of an example state are fully calculated. Suppose that the current state is *S*(31, 1) and that the template sequence is

TACCAAATGAGGTTATGGCTCTTTTTTGCATaagcattcctaaaaccatt

where the uppercase letters denote the first *l* = 31 positions which have already been transcribed. The next nucleotide to be incorporated onto the mRNA is therefore U.

Suppose the kinetic model is {𝕀𝕊 = 0, 𝔾𝕋 = 1, 𝕋𝕊 = *H*_*I*_*G*_*I*_, ℝ𝔹 = 1, 𝔹𝕋 = 1, ℍ𝕋 = 1}. Thus, the state pathway is

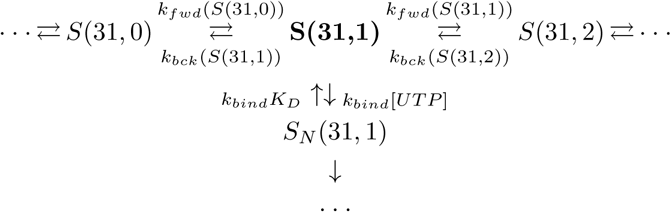

Assume that the transcription bubble is described by *h* = 11, *β*_1_ = 2, and *β*_2_ = 0. The baseline Gibbs energy barrier 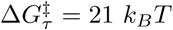 and the hypertranslocation barrier 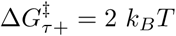. All other parameters used in this example are specified in Table 1.

In order to stochastically sample the next state (using the Gillespie algorithm [53, 55]), three rates must be calculated.

First, computing the rate of NTP binding is straightforward.

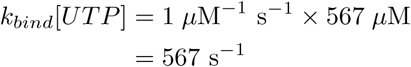

Second, because ℝ𝔹 = 1, in order to evaluate *k*_*bck*_(*S*(31, 1)) we must first compute the upstream mRNA secondary structure to check for steric collisions. Using Equation 2, the first 13 nucleotides of the mRNA strand are free to fold.

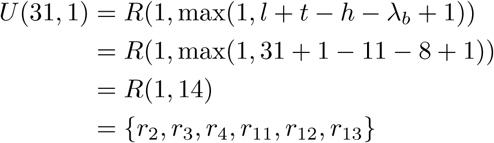

The final calculation in the above equation describes an RNA hairpin and was performed using the ViennaRNA suite [59]. Now, we can compute the Gibbs energy of the transition state for this backward translocation, and its associated rate constant

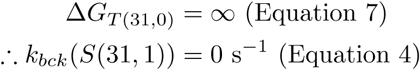

Backwards translocation is impossible due to steric collisions with the RNA hairpin (because of Equation 7, Case 1: ℝ𝔹 = 1 and *r*_13_ ∈ *U* (31, 1)).

Third, because ℝ𝔹 = 1, in order to evaluate *k*_*fwd*_(*S*(31, 1)) the downstream mRNA structure must be approximated. As the current state is not backtracked, there is no downstream mRNA free to fold.

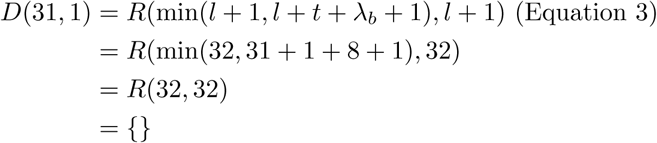

It is clear that Case 2 of Equation 7 will fail and Case 5 will instead apply (*t >* 0 and ℍ𝕋 = 1). Therefore, in order to calculate the rate of hypertranslocation, we must first compute the Gibbs energies of *S*(31, 1) and *T* (31, 1). The former term has Gibbs energy

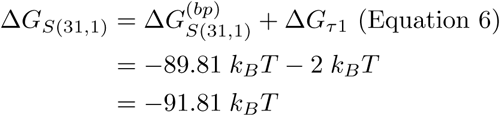

where 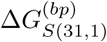 is computed using DNA/DNA and DNA/RNA thermodynamic parameters [56, 57], including the estimated dangling end contribution. According to the 𝕋𝕊 = *H*_*I*_*G*_*I*_ transition state model, the transition state is comprised of the basepairs which exist in both *S*(31, 1) and *S*(31, 2) (ie. the intersection). Given the values of *β*_1_, *β*_2_, and *h*, the DNA/DNA gene basepairs are

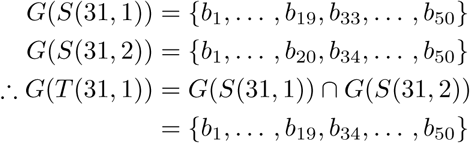

Similarly, the DNA/RNA hybrid basepairs are

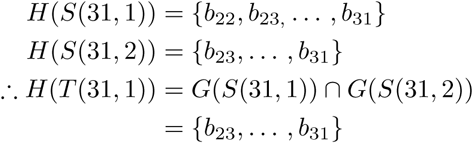

Therefore, by applying DNA/DNA parameters to *G*(*T* (31, 1)), the DNA/RNA parameters to *H*(*S*(31, 2)), and the normalisation constant of 𝕋𝕊 = *H*_*I*_*G*_*I*_, the Gibbs energy of the transition state is

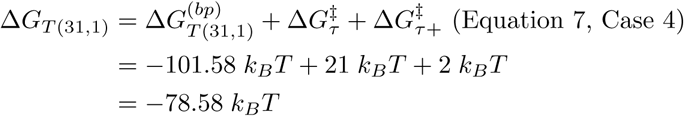

We can now calculate the forward translocation rate using Equation 5

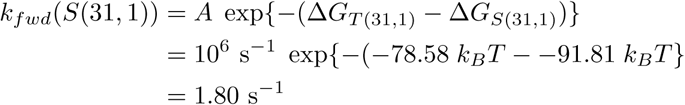

Therefore, the three rate constants leading out of *S*(31, 1) are

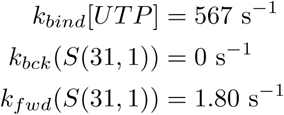

### Binary classification of pause sites

Pause site prediction is treated as a binary classification problem. Let *X* = (*X*_1_, *X*_2_, *…, X*_*L*_) be the nucleotide sequence of a gene of length *L*. Site *l* can be classified as a pause, *C*_*l*_ =𝒫, or as not being a pause, *C*_*l*_ = 𝒩.

Two binary classifiers are described. First, the kinetic model classifier (KMC) predicts pause sites based off the simulated time to catalysis of each genomic position. Second, the naive Bayes classifier (NBC) predicts pause sites using a simple nucleotide-windowed statistical technique.

19,960 pause sites from across the entire *E*. *coli* genome were identified in a high throughput *in vivo* analysis [28]. This dataset contains the “known” 𝒫 and 𝒩 classifications which are used to train and test the two classifiers. Using a full *E*. *coli* genome sequence (Accession: NC 000913), we randomly partitioned these genes into a training set and a test set. Due to computational limitations of the KMC, the two classifiers are trained on 50 genes and tested on the remaining 2403 genes. An additional 100 nt were included at the upstream and downstream ends of the translated region of each gene to account for boundary effects.

Classifiers are assessed by the area under the curve (AUC) of a receiver operating characteristic curve (ROC curve). The AUC quantifies the amount of information in a classifier and accounts for true positive and false positive rates simultaneously. AUC is bounded in the range [0, 1]. If the AUC is less than 0.5, then the classifier provides no information and is no better than random, while a perfect classifier would have an AUC of 1.0. See the review by Fawcett 2006 [62] for an introduction to ROC curves.

#### Kinetic model classifier

Classifying site *X*_*l*_ into class 𝒫 or 𝒩 is achieved by simulating the kinetic model using the Gillespie algorithm [53, 55]. Let *f*_𝒫_ (*l*) be the median time that the length of the mRNA is exactly *l* nucleotides in length.

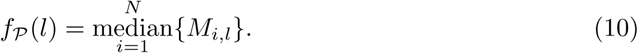

where *M* is an *N* × *L* matrix, where entry (*i, l*) is the total time that the mRNA contains exactly *l* nucleotides during simulation *i*, and *N* is the number of simulation trials performed on each gene (*N* = 100).

Site *l* is classified as a pause site if and only if *f*_𝒫_ (*l*) *> θ* for some threshold *θ*. Applying changes to the value of *θ* is required to build a ROC curve.

#### Naive Bayes classifier

The naive Bayes classifier is a simple probabilistic classifier derived from Bayes’ theorem [63] and makes a suitable bioinformatics algorithm for sequence-based prediction [64–66]. Classification of site *l* into 𝒫or 𝒩 is computed through comparison of the respective log posterior probabilities:

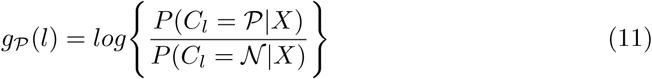

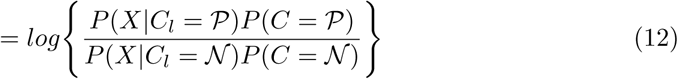

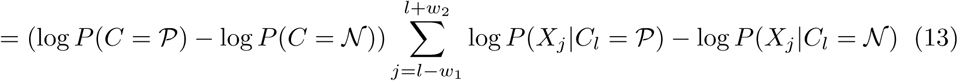

The naive property of the NBC invokes the assumption of independence between sites, allowing the likelihood to be computed as the product of likelihoods *P* (*X*_*j*_|*C*_*l*_ = *c*) across all sites in a window around *l*, where the window size is *w*_1_ + 1 + *w*_2_ (for *w*_1_ = 10 and *w*_2_ = 4). Log probabilities are trained using Laplace smoothing.

Site *l* is classified as a pause if *g*_*P*_ (*l*) > *θ* for some *θ*.

## Results

### Searching the space of kinetic models and parameters

Our aim was to 1) use Bayesian inference to select the best of 128 models (Fig 3); and 2) estimate the parameters. This was achieved using the rejection approximate Bayesian computation algorithm (R-ABC) [67, 68].

**Fig 3.**
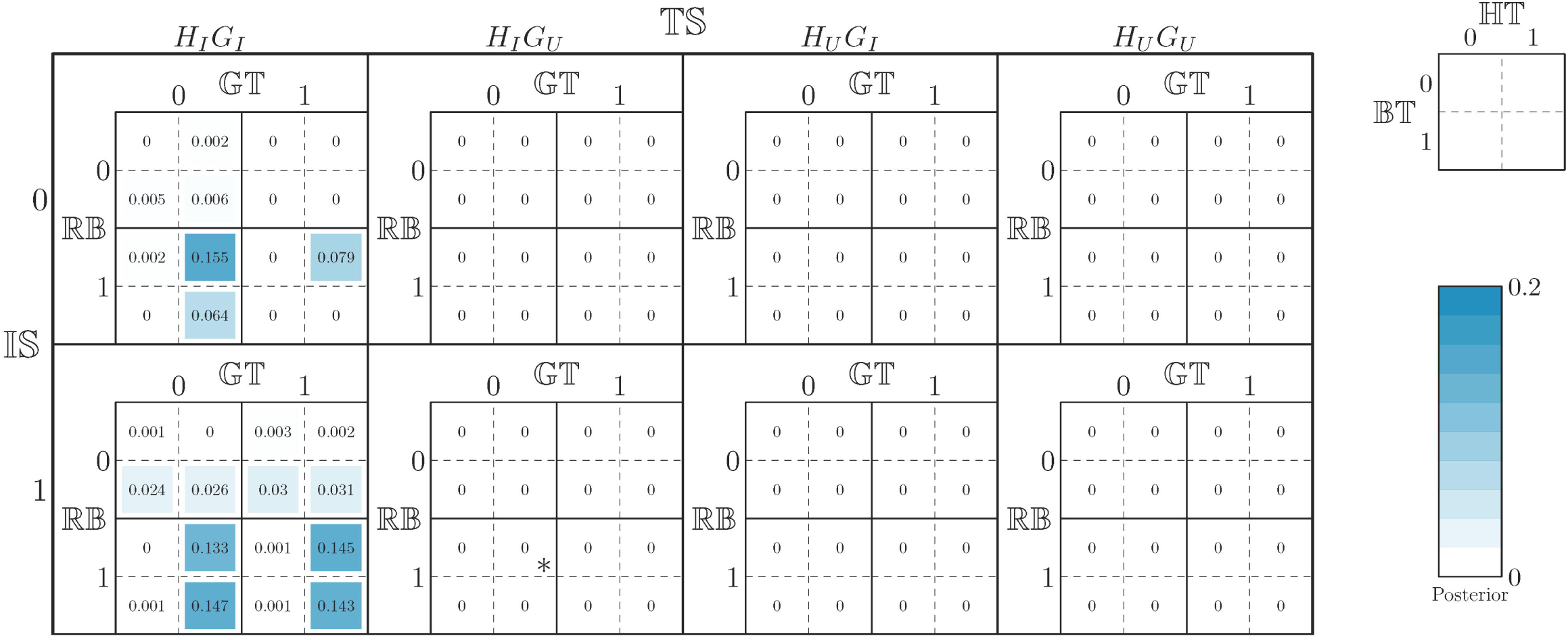
Comparison of kinetic models. Each square in the grid corresponds to one of the 128 kinetic models. For example, the grid square indicated with a star corresponds to the kinetic model which includes the intermediate state, does not model the gating tyrosine, uses the *H*_*I*_*G*_*U*_ model of translocation, employs the RNA blockade model, and includes hypertranslocation but not backtracking. This model is denoted by{𝕀𝕊 = 1, 𝔾𝕋 = 0, 𝕋𝕊 = *H*_*I*_*G*_*U*_, ℝ𝔹 = 1, ℍ𝕋 = 1, 𝔹𝕋 = 0}. Every square is shaded with opacity proportional to its kinetic model’s posterior probability and is labelled with this probability.

The kinetic-model-classifier is trained under a Bayesian framework: therefore the goal is not to optimise the AUC but rather sample parameters and models from the posterior distribution. However only parameters and models whose simulated value of 1-AUC is less than threshold *E* are accepted into the posterior distribution (S2 Appendix).

The results of this analysis are displayed in Fig 3, 4, and 5. These results show that the model with the highest posterior probability (of 0.155) is the model denoted by {𝕀𝕊 = 0, 𝔾𝕋 = 0, 𝕋𝕊 = *H*_*I*_*G*_*I*_, ℝ𝔹 = 1, ℍ𝕋 = 1, 𝔹𝕋 = 0}. The geometric median (ie. a point-estimate) of the posterior distribution has an AUC of 0.730 on the test set (and 0.746 on the training set), suggesting that while the kinetic model captures a reasonable amount of information about transcriptional pausing, it is still a fairly weak classifier.

**Fig 4.**
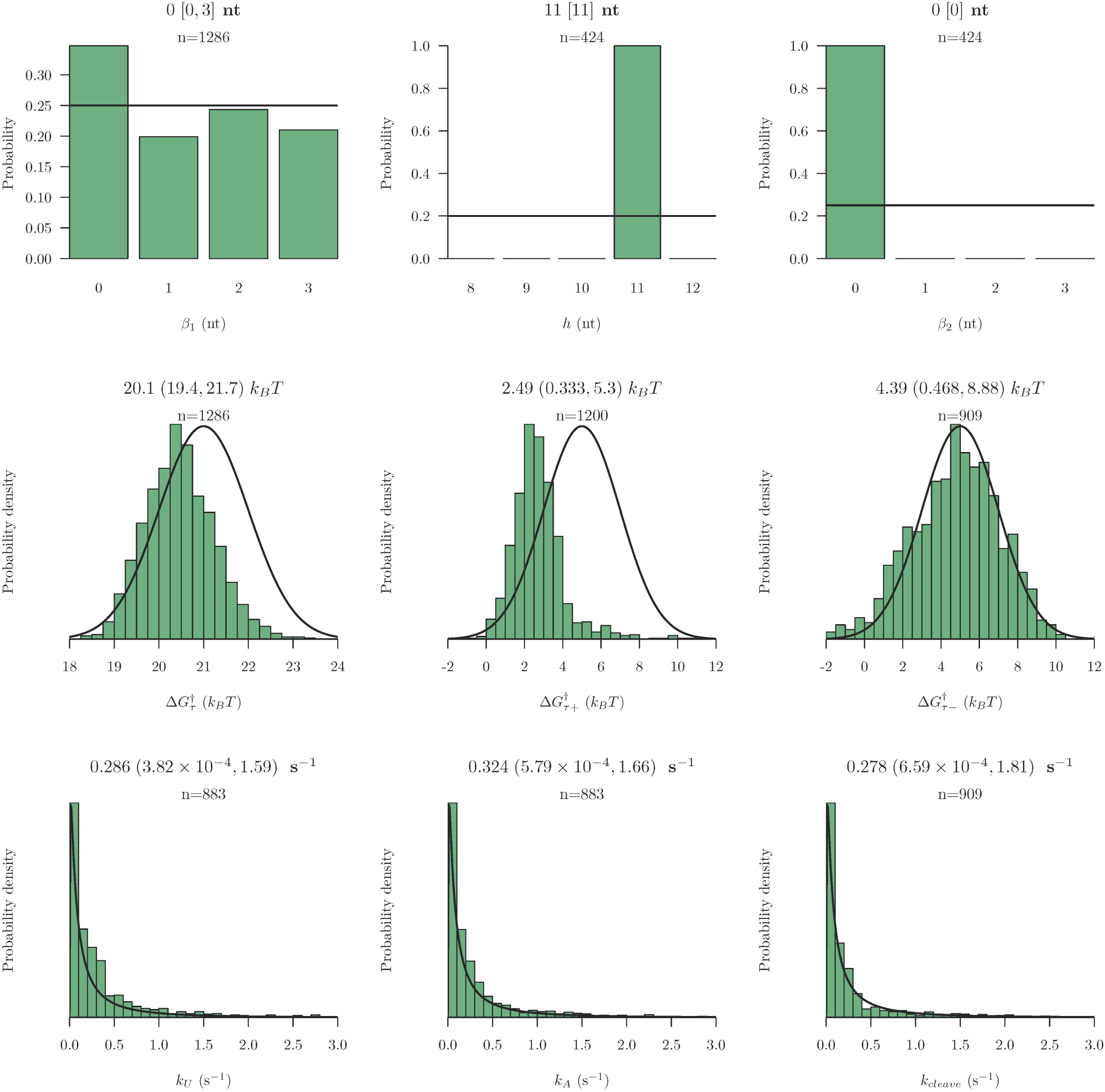
Posterior distribution of kinetic model parameters. Posterior distributions (coloured bars) and prior distributions (black curves) for the 9 estimated parameters are shown. Posterior distributions reported are conditional on the models which use the parameters. For example 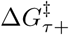 is conditional on ℍ𝕋 = 1, while *k*_*U*_ and *k*_*A*_ estimates are conditional on 𝕀𝕊 = 1. The geometric median point-estimate, the 95% highest posterior density (HPD) interval (calculated using Tracer 1.6 [69]), and posterior distribution sample size *n* are displayed above each plot (3 sf). These results reveal that *h, β*_2_, 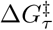, and 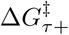 are informed by the pause site data, while the remaining parameters are largely uninformed.

**Fig 5.**
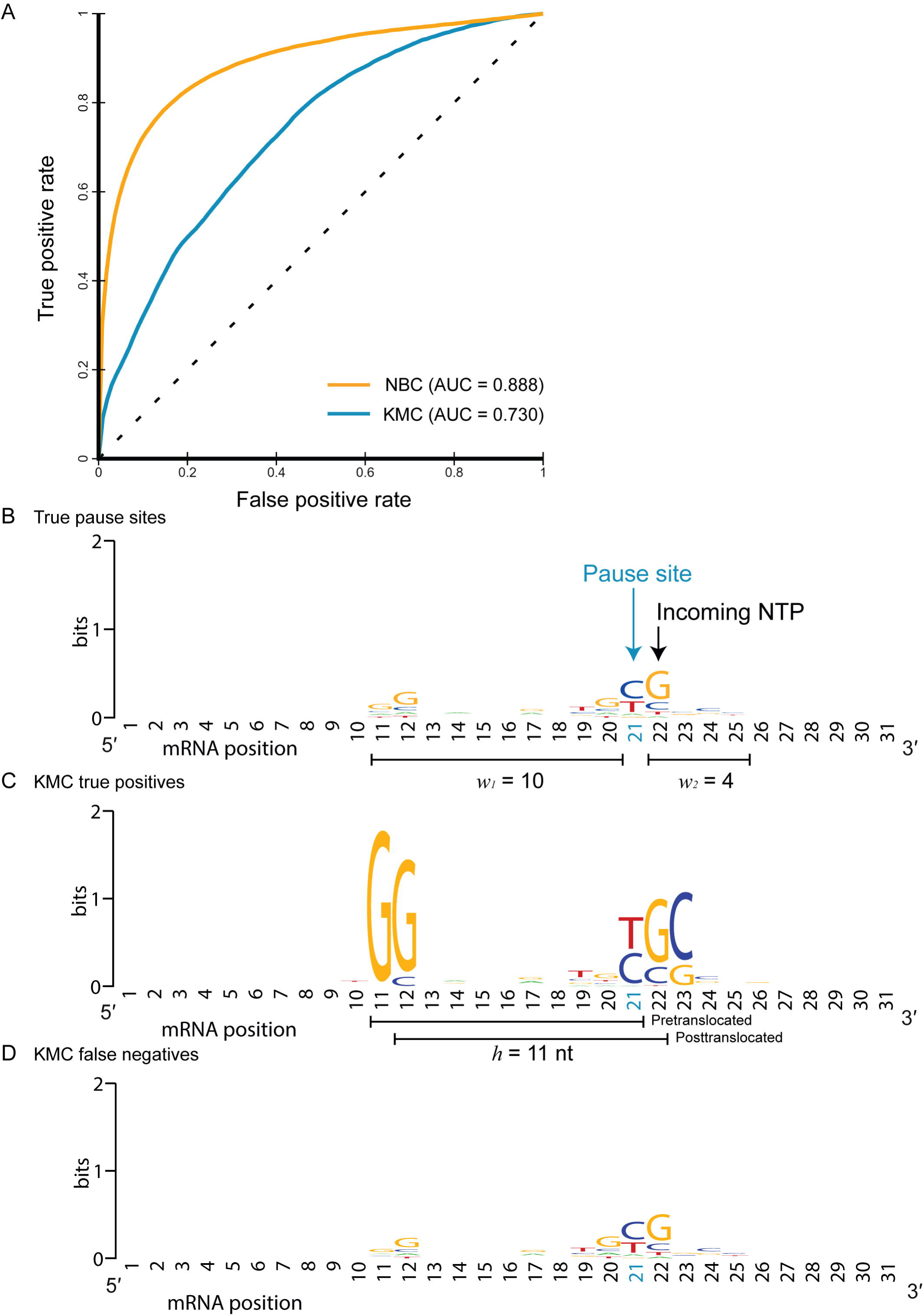
Classification of pause sites. A: A ROC curve comparing the predictive power of the kinetic model classifier (KMC) and the naive Bayes classifier (NBC). Sequence logos built using B: known pause sites, C: the subset of known pause sites which are correctly classified by the KMC (true positives), and D: the subset of known pause sites which are not classified as pause sites by the KMC (false negatives). The true positives and false negatives collectively comprise the true pause sites. The nucleotide window used by the NBC and the estimated hybrid length are displayed. All logos are generated using WebLogo [70] and trained on test set sequences.

### Assessment of feature importance

Our analysis suggests that 𝕋𝕊 is a critical model setting while *h* and *β*_2_ are critical parameters. There is evidence that ℍ𝕋 and ℝ𝔹 may also be critical. We wanted to confirm that these variables are indeed important for the prediction of pause sites, and to quantify how much predictive power is contained in each variable. To do this, we recomputed the AUC of models, by sampling from the posterior distribution, using samples that differ in the described variables (Table 2).

**Table 2.**
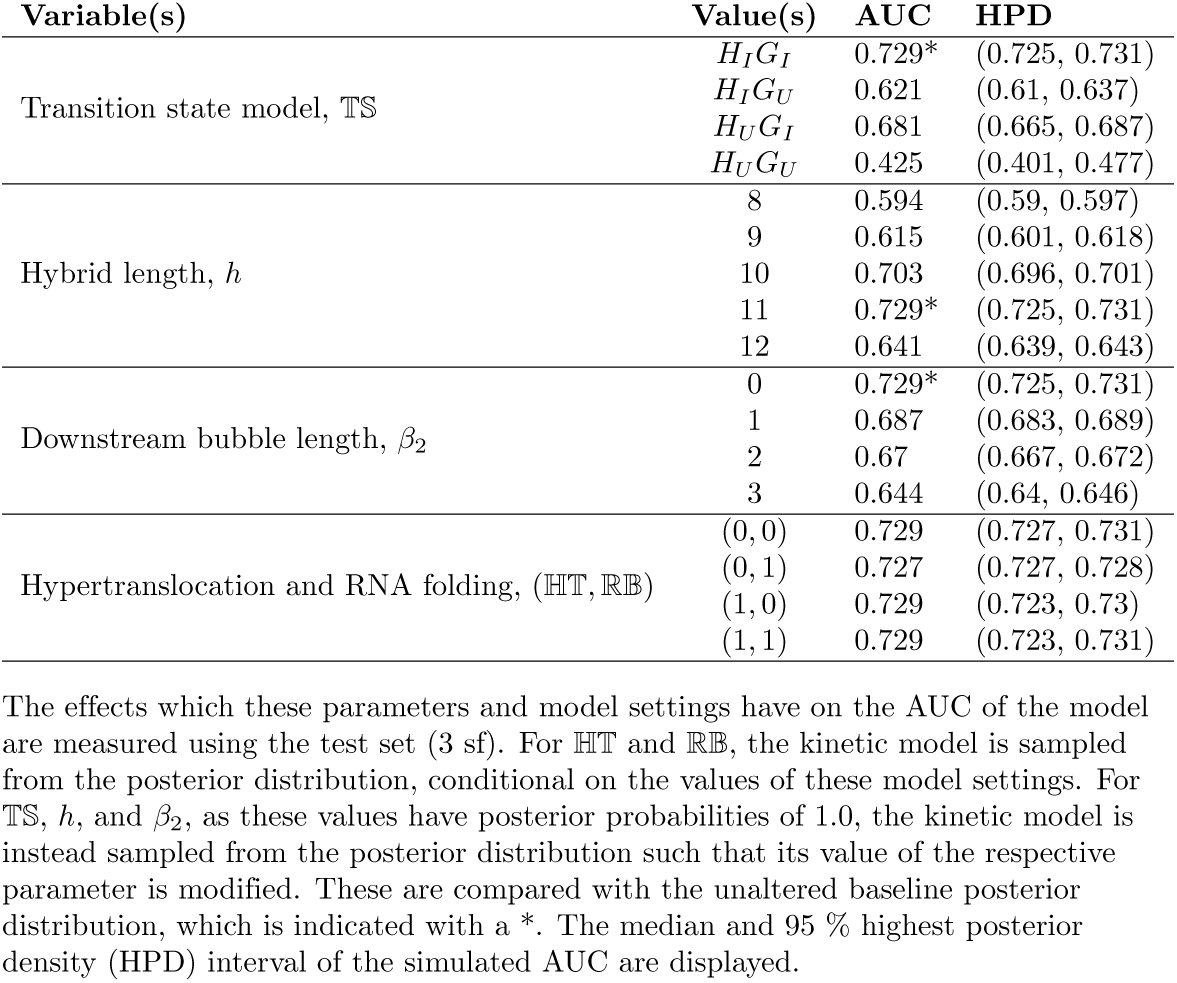
Effects of critical parameters and model settings on AUC.

| Variable(s) | Value(s) | AUC | HPD |
| --- | --- | --- | --- |
| Transition state model, TS | $H_I G_I$ | 0.729* | (0.725, 0.731) |
| | $H_I G_U$ | 0.621 | (0.61, 0.637) |
| | $H_U G_I$ | 0.681 | (0.665, 0.687) |
| | $H_U G_U$ | 0.425 | (0.401, 0.477) |
| Hybrid length, $h$ | 8 | 0.594 | (0.59, 0.597) |
|  | 9 | 0.615 | (0.601, 0.618) |
|  | 10 | 0.703 | (0.696, 0.701) |
|  | 11 | 0.729* | (0.725, 0.731) |
|  | 12 | 0.641 | (0.639, 0.643) |
| Downstream bubble length, $\beta_2$ | 0 | 0.729* | (0.725, 0.731) |
|  | 1 | 0.687 | (0.683, 0.689) |
|  | 2 | 0.67 | (0.667, 0.672) |
|  | 3 | 0.644 | (0.64, 0.646) |
| Hypertranslocation and RNA folding, (HT, RB) | (0, 0) | 0.729 | (0.727, 0.731) |
|  | (0, 1) | 0.727 | (0.727, 0.728) |
|  | (1, 0) | 0.729 | (0.723, 0.73) |
|  | (1, 1) | 0.729 | (0.723, 0.731) |

The effects which these parameters and model settings have on the AUC of the model are measured using the test set (3 sf). For HT and RB, the kinetic model is sampled from the posterior distribution, conditional on the values of these model settings. For TS, $h$ , and $\beta_2$ , as these values have posterior probabilities of 1.0, the kinetic model is instead sampled from the posterior distribution such that its value of the respective parameter is modified. These are compared with the unaltered baseline posterior distribution, which is indicated with a \*. The median and 95 % highest posterior density (HPD) interval of the simulated AUC are displayed.

These results confirm that 𝕋𝕊, *h*, and *β*_2_ are indeed critical variables. In the most extreme case, changing the transition state model 𝕋𝕊 from *H*_*I*_*G*_*I*_ to *H*_*U*_ *G*_*U*_ reduced the AUC from 0.73 to 0.43; the latter corresponding to a predictive model that is worse than assigning pause sites at random. In the least extreme case, changing *h* from 11 nt to 10 nt reduced the AUC from 0.73 to 0.70.

Whereas, adjusting sampling from the posterior distribution, conditional on ℍ𝕋 and ℝ𝔹, did not yield any significant AUC changes across the four pairwise combinations of ℍ𝕋 and ℝ𝔹 (other than a minor decrease in AUC for {ℍ𝕋 = 0, ℝ𝔹 = 1}). It is therefore likely that these two model settings are not offering any further predictive power.

### Naive Bayes model

We trained a naive Bayes classifier to predict pause sites using the same dataset [28] as the kinetic model. This model enabled an estimation of the amount of information available in the data, without the constraint of being physically plausible.

Similar to the kinetic model, we performed a ROC analysis on the NBC to evaluate how accurately it can predict pause sites (Fig 5). The AUC of the NBC was 0.888 on the test set (and 0.895 on training set), suggesting that the sequence within this window contains a large amount of information about transcriptional pausing, even though the sites are assessed independently. This model has significantly better prediction power than the kinetic model. The extent of overfitting is minimal.

## Discussion

In this study we inferred structural, kinetic, and thermodynamic parameters (Fig 4) of kinetic models for the prediction of transcriptional pausing. We also inferred the kinetic model itself to provide evidence for or against various kinetic model variants that have been described in the literature (Fig 3). The posterior distribution of kinetic models is interpreted using the Bayes factor, *K*, following the general guidelines by Kass and Raftery 1995 [71]. We compared the predictive power of the kinetic models to that of a statistical technique: the naive Bayes classifier (Fig 5).

These results suggest that the translocation transition model 𝕋𝕊 and structural parameters governing the transcription bubble (*h* and *β*_2_) are heavily informed by the pause site data and are important parameters for the prediction of pause sites. The relative Gibbs energies of the pre and posttranslocated state, and the rate of translocation between the two, is the primary effector of pausing. In contrast, the data tells us little about the remaining parameters and model settings. Overall, the kinetic model contains a moderate degree of predictive power (AUC = 0.73), but is severely lacking compared with the NBC (AUC = 0.89).

### The structure of the transcription bubble strongly affects pausing

We estimate that the transcription bubble contains an *ĥ* = 11 nt hybrid and a 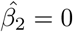 nt gap between the RNAP and the downstream gene region (Fig 4). These estimates each have a posterior probability of 1.00, indicative of high certainty that these are the best estimates for the data. In contrast the gap between the RNAP and the upstream dsDNA 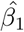 was not informed by the data.

Furthermore, the translocation transition model 𝕋𝕊 is an extremely important model setting. We estimate that the best transition model estimate is 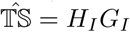 (Fig 3). This variant has a marginal posterior probability *P* (𝕋𝕊 = *H*_*I*_*G*_*I*_|*D*) = 1.0, indicating high confidence that this is the preferred transition state model. Our critical parameter analysis reveals that by changing the value of 𝕋𝕊, while holding all else constant, in the extreme case the AUC declines from 0.73 down to 0.48 (Table 2).

Under the *H*_*I*_*G*_*I*_ model, the transition state between two translocation states contains the intersection between the two sets of DNA/RNA hybrid basepairs, and the intersection between the two sets of DNA/DNA gene basepairs.

This model corresponds to a physical mechanism of translocation akin to a bubble melting process (see Fig 2A). First, one DNA/DNA basepair within the gene and one DNA/RNA basepair within the hybrid break, eg. due to thermal collisions. These two bond breaking events could happen in any order and facilitate the formation of the translocation transition state. Assuming that *β*_1_ = 0, *h* = 11, and *β*_2_ = 0 in a regular translocation state, the effective values of these parameters would be *β*_1_ = 1, *h* = 10, and *β*_2_ = 1 in the transition state. The transcription bubble is therefore one nucleotide wider (from 11 to 12 nucleotides). Second, RNAP is able to diffuse into this opening and, in no specific order, one basepair within the gene and one basepair within the hybrid form. This completes the translocation step.

Examining the sequence logo generated from known pause sites (Fig 5B), we can see that the nucleotides 1-2 positions proximal to the pause site are important, as are the two nucleotides 9-10 positions upstream from the pause site. Pausing usually occurs at a cytidine or uridine and the incoming NTP is usually guanosine. Comparing this with the sequence logo generated from the pause sites which were correctly classified by the kinetic model (Fig 5C), we can see that some of this pattern is captured by the kinetic model. The two nucleotides 9-10 positions upstream from the pause site exist at the upstream end of the *h* = 11 nt DNA/RNA hybrid, and the two nucleotides 1-2 positions downstream from the pause site exist *β*_2_ = 0 nt downstream of the hybrid within the DNA/DNA gene. These two regions constitute two basepair doublets and, under the 𝕋𝕊 = *H*_*I*_*G*_*I*_ model, must be broken to facilitate forward translocation into the transition state and therefore into the posttranslocated position. These two doublets tend to be G-C rich, which, under the nearest neighbour models, correspond to stronger basepairs that require more energy to break than A-U or A-T basepairs [56, 57].

However, the kinetic model is unable to explain pause sites which do not have strong basepairs at the upstream end of the hybrid. The kinetic model also places too much weight on the position two nucleotides downstream from the pause site (position 23 in the sequence logos). The reasonably large differences in positional information between these two sequence logos suggests that the kinetic model is unable to model pausing when the sequence composition even slightly deviates from this motif.

Estimates of *β*_1_ and *β*_2_ are consistent with previous estimates from crystal structures [4–7]. However, based off these same structures the pretranslocated hybrid length is estimated as 10 bp, which is inconsistent with our estimate of *ĥ* = 11 bp. This is a peculiar contradiction, especially considering that our Bayesian protocol is 100% certain that *ĥ* = 11 bp, so it may be more suitable to consider *h* not as being the true hybrid size but rather the *effective* hybrid size during transcription.

Overall, there is strong evidence that by having accurate estimates for the parameters which govern the size of the transcription bubble, and a good model of the translocation transition state, the sequence-specific properties that emerge are important for the prediction of pause sites. If two RNA polymerases were to have different transcription bubble dynamics, they would behave quite differently on the same sequence.

### Backtracking and the intermediate state are not necessary to predict pausing

By combining three model variants – 𝔹𝕋, 𝔾𝕋, and 𝕀𝕊 – in a combinatorial fashion, eight plausible variants of transcriptional pausing, through off-pathway upstream events, were compared. Backtracking models were examined, and the Gibbs energy barrier of backtracking 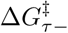 and the rate of RNA cleavage *k*_*cleave*_ were estimated. The hypothesised role the gating tyrosine plays in eliciting pauses, by permitting rapid translocation into *S*(*l*, − 1) while delimiting further backstepping, was incorporated with the backtracking model (Fig 2D). Catalytically inactive intermediate state models were assessed and the rates of entry *k*_*U*_ and exit *k*_*A*_ to and from this state were estimated.

And yet, none of these eight variants were notably superior in their ability to predict the locations of pause sites. These three mechanisms have marginal posterior probabilities ranging from 0.44 to 0.69 (Fig 3), to be interpreted as ‘not worth more than a bare mention’ [71]. The four related parameters – 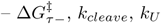, and *k*_*A*_ – have posterior distributions almost the same as their prior distributions, consistent with the data providing no information about these parameters.

A posterior probability of intermediate magnitude is indicative of the model setting being neither necessary nor detrimental. Instead the feature is unnecessary.

Our findings are consistent with the hypothesis that the relative stability between the pre and posttranslocated states chiefly facilitates the occurrence of pausing [21, 51], as opposed to the stability of those relative to the backtracked states [18, 44, 72]. Rather, most pauses are brief (averaging 3 s) and do not involve backtracking [42]. It is likely that backtracking occurs on a timescale so slow [73] that by the time RNAP has had sufficient time to sample the energy landscape of backtracked states, the pause has already begun, and therefore these energies are of little use for predicting the frequency of pausing.

Pauses are hypothesised to occur largely through the conformational rearrangement of the enzyme into a catalytically inactive form – the IS [1, 18–22]. This may occur in a sequence-dependent manner where weaker DNA/RNA hybrids are more likely to invoke this transition [18, 19, 34]. The sequence-independent model of this transition used in this study was also of no use for predicting the locations of pause sites.

Although, backtracking, the gating tyrosine, and the IS were not able to explain the *frequency* of pausing, they may be able to explain the *duration* of pausing. Treating this system as a regression problem – by fitting to known pause *durations* – as opposed to a classification problem – where pausing is viewed as a binary trait – may offer further insights into these model features.

### Hypertranslocation and mRNA folding are not necessary to predict pausing

Our initial analysis suggested that hypertranslocation ℍ𝕋 could be a necessary model feature; with marginal posterior probability *P* (ℍ𝕋 = 1*|D*) = 0.93 – corresponding to a Bayes factors of 13.3 – therefore providing ‘positive evidence’ that 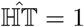 [71].

Accordingly, the posterior distribution of hypertranslocation Gibbs barrier 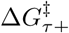 is quite different to its prior distribution (Fig 4).

There was also evidence that incorporating RNA folding into the model may be necessary; *P* (ℝ𝔹 = 1|*D*) = 0.87 and a Bayes factor of 6.7; ‘positive evidence’ [71]. In the ℝ𝔹 = 1 model, upstream (downstream) RNA secondary structures inhibit backward (forward) translocation.

However, when these two model settings were systematically evaluated on the test set, they were not found to be necessary. Subsampling from the posterior distribution conditional on ℍ𝕋 and ℝ𝔹 did not yield any significant change in AUC across the four pairwise combinations of ℍ𝕋 and ℝ𝔹, aside from a trivial decrease in AUC for {ℍ𝕋 = 0, ℝ𝔹 = 1 (Table 2). It is therefore likely that although ℍ𝕋 = 1 and ℝ𝔹 = 1} are associated with higher posterior probabilities, the model which has these settings enabled {ℍ𝕋 = 1, ℝ𝔹 = 1} does not contain any predictive power beyond that of {ℍ𝕋 = 0, ℝ𝔹 = 0}, so the hypertranslocation and RNA blockade models are of little to no use for the prediction of pause sites.

These results are consistent with the findings of Levint et al. 1987, who found no correlation between the locations of pause sites and upstream RNA secondary structures [74], and Dalal et al. 2006 [75], who used optical tweezers to inhibit mRNA folding during transcription, and found that the kinetics of pausing were not affected by the perturbation.

This is not to say that these mechanisms are not involved in pausing. RNA folding and hypertranslocation cooperatively induce pausing at the *his* leader pause site (a Class I pause), for instance. This is achieved by direct interaction between an upstream hairpin and the RNAP [18]. However, on average, the described models of RNA folding and hypertranslocation are of no assistance for the prediction of pause sites.

### Predicting and explaining transcriptional pausing

The transcription kinetic model successfully predicted pause sites (and non-pause sites) to a moderate level of accuracy (AUC = 0.73). The sequence-dependence of the kinetic model emerges primarily from the Gibbs energies of basepairing. Therefore the AUC is approximately a measure of how much information basepairing thermodynamics have about transcriptional pause sites.

However, the NBC was significantly more successful at predicting the locations of pause sites (AUC = 0.89). While there may exist other machine learning techniques that perform even better than the NBC [50], this provides a lower-bound of the amount of information contained in the data. With a sufficient understanding of the kinetics of transcription, the kinetic model classifier should perform at least as well as the NBC did.

Gibbs energies of basepairing can only take the kinetic model so far. Physically informed features that could extract the full potential of the kinetic model classifier include: the effects of double-stranded DNA bending upstream and downstream of the polymerase [19, 43]; a sequence-dependent model of entry into the IS [18, 19, 34]; specific interactions between the DNA/RNA hybrid and the protein [22]; effects that the promoter have on the way the polymerase interacts with the gene [76]; differential rates of NTP binding, release, and catalysis across the four nucleotides [46]; effects that the nucleotide context around the 3′ mRNA have on the rates of NTP incorporation [77]; or the effects of local NTP depletion [78]. A mathematical understanding of such processes, among others, may be necessary for the kinetic model to perform as well as a sequence motif and bridge the gap between the two sequence logos (Fig 5).

## Conclusion

In this study, we quantified the predictive power that standard kinetic models of transcription elongation have with respect to identifying transcriptional pause sites. Transcriptional pausing is not sufficiently understood to capture the signal from the data as thoroughly as a non-physical statistical model. We suggest that the relative stability between the pre and posttranslocated states, and the estimated energy required to translocate between them, is the primary effector of pausing. Backtracking, hypertranslocation, RNA blockades, and the catalytically inactive intermediate state are not required to explain whether pausing occurs, however they may explain pause duration.

## Supporting information

**S1 Appendix**. **Prior distribution justifications**. Brief justifications for the prior distributions presented in Table 1.

**S2 Appendix**. **Approximate Bayesian Computation**. A description of the R-ABC algorithm for the inference of parameters and models.

## References

1. Maoiléidigh DÓ, Tadigotla VR, Nudler E, Ruckenstein AE. A unified model of transcription elongation: what have we learned from single-molecule experiments? Biophysical journal. 2011;100(5):1157–1166.

2. Nudler E, Mustaev A, Goldfarb A, Lukhtanov E. The RNA–DNA hybrid maintains the register of transcription by preventing backtracking of RNA polymerase. Cell. 1997;89(1):33–41.

3. Gnatt AL, Cramer P, Fu J, Bushnell DA, Kornberg RD. Structural basis of transcription: an RNA polymerase II elongation complex at 3.3 °A resolution. Science. 2001;292(5523):1876–1882.

4. Vassylyev DG, Vassylyeva MN, Perederina A, Tahirov TH, Artsimovitch I. Structural basis for transcription elongation by bacterial RNA polymerase. Nature. 2007;448(7150):157.

5. Guo X, Myasnikov AG, Chen J, Crucifix C, Papai G, Takacs M, et al. Structural basis for NusA stabilized transcriptional pausing. Molecular cell. 2018;69(5):816–827.

6. Kang JY, Olinares PDB, Chen J, Campbell EA, Mustaev A, Chait BT, et al. Structural basis of transcription arrest by coliphage HK022 Nun in an Escherichia coli RNA polymerase elongation complex. Elife. 2017;6:e25478.

7. Vassylyev DG, Sekine Si, Laptenko O, Lee J, Vassylyeva MN, Borukhov S, et al. Crystal structure of a bacterial RNA polymerase holoenzyme at 2.6 °A resolution. Nature. 2002;417(6890):712.

8. Bar-Nahum G, Epshtein V, Ruckenstein AE, Rafikov R, Mustaev A, Nudler E. A ratchet mechanism of transcription elongation and its control. Cell. 2005;120(2):183–193.

9. Abbondanzieri EA, Greenleaf WJ, Shaevitz JW, Landick R, Block SM. Direct observation of base-pair stepping by RNA polymerase. Nature. 2005;438(7067):460–465.

10. Komissarova N, Kashlev M. Transcriptional arrest: Escherichia coli RNA polymerase translocates backward, leaving the 3/end of the RNA intact and extruded. Proceedings of the National Academy of Sciences. 1997;94(5):1755–1760.

11. Cheung AC, Cramer P. Structural basis of RNA polymerase II backtracking, arrest and reactivation. Nature. 2011;471(7337):249.

12. Toulmé F, Mosrin-Huaman C, Sparkowski J, Das A, Leng M, Rahmouni AR. GreA and GreB proteins revive backtracked RNA polymerase in vivo by promoting transcript trimming. The EMBO journal. 2000;19(24):6853–6859.

13. Lisica A, Engel C, Jahnel M, Roldán É, Galburt EA, Cramer P, et al. Mechanisms of backtrack recovery by RNA polymerases I and II. Proceedings of the National Academy of Sciences. 2016; p. 201517011.

14. Yarnell W, Roberts J. Mechanism of intrinsic transcription termination and antitermination. Science. 1999;284(5414):611–615.

15. Larson MH, Greenleaf WJ, Landick R, Block SM. Applied force reveals mechanistic and energetic details of transcription termination. Cell. 2008;132(6):971–982.

16. Santangelo TJ, Roberts JW. Forward translocation is the natural pathway of RNA release at an intrinsic terminator. Molecular cell. 2004;14(1):117–126.

17. Gusarov I, Nudler E. The mechanism of intrinsic transcription termination. Molecular cell. 1999;3(4):495–504.

18. Artsimovitch I, Landick R. Pausing by bacterial RNA polymerase is mediated by mechanistically distinct classes of signals. Proceedings of the National Academy of Sciences. 2000;97(13):7090–7095.

19. Palangat M, Hittinger CT, Landick R. Downstream DNA selectively affects a paused conformation of human RNA polymerase II. Journal of molecular biology. 2004;341(2):429–442.

20. Herbert KM, Greenleaf WJ, Block SM. Single-molecule studies of RNA polymerase: motoring along. Annu Rev Biochem. 2008;77:149–176.

21. Imashimizu M, Kireeva ML, Lubkowska L, Gotte D, Parks AR, Strathern JN, et al. Intrinsic translocation barrier as an initial step in pausing by RNA polymerase II. Journal of molecular biology. 2013;425(4):697–712.

22. Saba J, Chua X, Mishanina TV, Nayak D, Windgassen TA, Mooney RA, et al. The elemental mechanism of transcriptional pausing. bioRxiv. 2018; p. 422220.

23. Herbert KM, La Porta A, Wong BJ, Mooney RA, Neuman KC, Landick R, et al. Sequence-resolved detection of pausing by single RNA polymerase molecules. Cell. 2006;125(6):1083–1094.

24. Toulokhonov I, Zhang J, Palangat M, Landick R. A central role of the RNA polymerase trigger loop in active-site rearrangement during transcriptional pausing. Molecular cell. 2007;27(3):406–419.

25. Ryals J, Little R, Bremer H. Temperature dependence of RNA synthesis parameters in Escherichia coli. Journal of bacteriology. 1982;151(2):879–887.

26. Vogel U, Jensen KF. The RNA chain elongation rate in Escherichia coli depends on the growth rate. Journal of bacteriology. 1994;176(10):2807–2813.

27. Mason PB, Struhl K. Distinction and relationship between elongation rate and processivity of RNA polymerase II in vivo. Molecular cell. 2005;17(6):831–840.

28. Larson MH, Mooney RA, Peters JM, Windgassen T, Nayak D, Gross CA, et al. A pause sequence enriched at translation start sites drives transcription dynamics in vivo. Science. 2014;344(6187):1042–1047.

29. Kireeva ML, Kashlev M. Mechanism of sequence-specific pausing of bacterial RNA polymerase. Proceedings of the National Academy of Sciences. 2009;106(22):8900–8905.

30. Shundrovsky A, Santangelo TJ, Roberts JW, Wang MD. A single-molecule technique to study sequence-dependent transcription pausing. Biophysical journal. 2004;87(6):3945–3953.

31. Klumpp S, Hwa T. Stochasticity and traffic jams in the transcription of ribosomal RNA: Intriguing role of termination and antitermination. Proceedings of the National Academy of Sciences. 2008;105(47):18159–18164.

32. Davis L, Gedeon T, Gedeon J, Thorenson J. A traffic flow model for bio-polymerization processes. Journal of mathematical biology. 2014;68(3):667–700.

33. Landick R. The regulatory roles and mechanism of transcriptional pausing; 2006.

34. Palangat M, Landick R. Roles of RNA: DNA hybrid stability, RNA structure, and active site conformation in pausing by human RNA polymerase II1. Journal of molecular biology. 2001;311(2):265–282.

35. Feng S, Holland EC. HIV-1 tat trans-activation requires the loop sequence within tar. Nature. 1988;334(6178):165.

36. Winkler WC, Cohen-Chalamish S, Breaker RR. An mRNA structure that controls gene expression by binding FMN. Proceedings of the National Academy of Sciences. 2002;99(25):15908–15913.

37. Wickiser JK, Winkler WC, Breaker RR, Crothers DM. The speed of RNA transcription and metabolite binding kinetics operate an FMN riboswitch. Molecular cell. 2005;18(1):49–60.

38. Oesterreich FC, Preibisch S, Neugebauer KM. Global analysis of nascent RNA reveals transcriptional pausing in terminal exons. Molecular cell. 2010;40(4):571–581.

39. Kornblihtt AR, de la Mata M, Fededa JP, Munoz MJ, Nogues G. Multiple links between transcription and splicing. Rna. 2004;10(10):1489–1498.

40. Roberts GC, Gooding C, Mak HY, Smith CW, Proudfoot NJ. Co-transcriptional commitment to alternative splice site selection. Nucleic acids research. 1998;26(24):5568–5572.

41. de la Mata M, Alonso CR, Kadener S, Fededa JP, Blaustein M, Pelisch F, et al. A slow RNA polymerase II affects alternative splicing in vivo. Molecular cell. 2003;12(2):525–532.

42. Neuman KC, Abbondanzieri EA, Landick R, Gelles J, Block SM. Ubiquitous transcriptional pausing is independent of RNA polymerase backtracking. Cell. 2003;115(4):437–447.

43. Kerppola TK, Kane CM. Analysis of the signals for transcription termination by purified RNA polymerase II. Biochemistry. 1990;29(1):269–278.

44. Tadigotla VR, Maoiléidigh DÓ, Sengupta AM, Epshtein V, Ebright RH, Nudler E, et al. Thermodynamic and kinetic modeling of transcriptional pausing. Proceedings of the National Academy of Sciences of the United States of America. 2006;103(12):4439–4444.

45. Hawryluk PJ, Újvári A, Luse DS. Characterization of a novel RNA polymerase II arrest site which lacks a weak 3’ RNA–DNA hybrid. Nucleic acids research. 2004;32(6):1904–1916.

46. Bai L, Fulbright RM, Wang MD. Mechanochemical kinetics of transcription elongation. Physical review letters. 2007;98(6):068103.

47. James K, Gamba P, Cockell SJ, Zenkin N. Misincorporation by RNA polymerase is a major source of transcription pausing in vivo. Nucleic acids research. 2016;45(3):1105–1113.

48. Imashimizu M, Oshima T, Lubkowska L, Kashlev M. Direct assessment of transcription fidelity by high-resolution RNA sequencing. Nucleic acids research. 2013;41(19):9090–9104.

49. Sydow JF, Brueckner F, Cheung AC, Damsma GE, Dengl S, Lehmann E, et al. Structural basis of transcription: mismatch-specific fidelity mechanisms and paused RNA polymerase II with frayed RNA. Molecular cell. 2009;34(6):710–721.

50. Siebert M, Söding J. Bayesian Markov models consistently outperform PWMs at predicting motifs in nucleotide sequences. Nucleic acids research. 2016;44(13):6055–6069.

51. Bai L, Shundrovsky A, Wang MD. Sequence-dependent kinetic model for transcription elongation by RNA polymerase. Journal of molecular biology. 2004;344(2):335–349.

52. Bochkareva A, Yuzenkova Y, Tadigotla VR, Zenkin N. Factor-independent transcription pausing caused by recognition of the RNA–DNA hybrid sequence. The EMBO journal. 2012;31(3):630–639.

53. Gillespie DT. Exact stochastic simulation of coupled chemical reactions. The journal of physical chemistry. 1977;81(25):2340–2361.

54. Lecca P. Stochastic chemical kinetics. Biophysical reviews. 2013;5(4):323–345.

55. Douglas J, Kingston R, Drummond A. Bayesian inference and comparison of stochastic transcription elongation models. bioRxiv. 2018; p. 499277.

56. SantaLucia J. A unified view of polymer, dumbbell, and oligonucleotide DNA nearest-neighbor thermodynamics. Proceedings of the National Academy of Sciences. 1998;95(4):1460–1465.

57. Wu P, Nakano Si, Sugimoto N. Temperature dependence of thermodynamic properties for DNA/DNA and RNA/DNA duplex formation. The FEBS Journal. 2002;269(12):2821–2830.

58. Flamm C, Fontana W, Hofacker IL, Schuster P. RNA folding at elementary step resolution. Rna. 2000;6(3):325–338.

59. Lorenz R, Bernhart SH, Zu Siederdissen CH, Tafer H, Flamm C, Stadler PF, et al. ViennaRNA Package 2.0. Algorithms for Molecular Biology. 2011;6(1):1.

60. Xia T, SantaLucia Jr J, Burkard ME, Kierzek R, Schroeder SJ, Jiao X, et al. Thermodynamic parameters for an expanded nearest-neighbor model for formation of RNA duplexes with Watson-Crick base pairs. Biochemistry. 1998;37(42):14719–14735.

61. Dangkulwanich M, Ishibashi T, Liu S, Kireeva ML, Lubkowska L, Kashlev M, et al. Complete dissection of transcription elongation reveals slow translocation of RNA polymerase II in a linear ratchet mechanism. Elife. 2013;2:e00971.

62. Fawcett T. An introduction to ROC analysis. Pattern recognition letters. 2006;27(8):861–874.

63. Kotsiantis SB, Zaharakis I, Pintelas P. Supervised machine learning: A review of classification techniques. Emerging artificial intelligence applications in computer engineering. 2007;160:3–24.

64. Cao J, Panetta R, Yue S, Steyaert A, Young-Bellido M, Ahmad S. A naive Bayes model to predict coupling between seven transmembrane domain receptors and G-proteins. Bioinformatics. 2003;19(2):234–240.

65. Rani P, Pudi V. RBNBC: Repeat based näive Bayes classifier for biological sequences. In: Data Mining, 2008. ICDM’08. Eighth IEEE International Conference on. IEEE; 2008. p. 989–994.

66. Feng PM, Ding H, Chen W, Lin H. Naive Bayes classifier with feature selection to identify phage virion proteins. Computational and mathematical methods in medicine. 2013;2013.

67. Beaumont MA. Approximate Bayesian computation in evolution and ecology. Annual review of ecology, evolution, and systematics. 2010;41:379–406.

68. Csilléry K, Blum MG, Gaggiotti OE, François O. Approximate Bayesian computation (ABC) in practice. Trends in ecology & evolution. 2010;25(7):410–418.

69. Rambaut A, Drummond A. Tracer 1.6. University of Edinburgh, Edinburgh. UK. Technical report; 2013.

70. Crooks GE, Hon G, Chandonia JM, Brenner SE. WebLogo: a sequence logo generator. Genome research. 2004;14(6):1188–1190.

71. Kass RE, Raftery AE. Bayes factors. Journal of the american statistical association. 1995;90(430):773–795.

72. Guajardo R, Sousa R. A model for the mechanism of polymerase translocation. Journal of molecular biology. 1997;265(1):8–19.

73. Galburt EA, Grill SW, Wiedmann A, Lubkowska L, Choy J, Nogales E, et al. Backtracking determines the force sensitivity of RNAP II in a factor-dependent manner. Nature. 2007;446(7137):820–823.

74. Levint JR, Chamberlin MJ. Mapping and characterization of transcriptional pause sites in the early genetic region of bacteriophage T7. Journal of molecular biology. 1987;196(1):61–84.

75. Dalal RV, Larson MH, Neuman KC, Gelles J, Landick R, Block SM. Pulling on the nascent RNA during transcription does not alter kinetics of elongation or ubiquitous pausing. Molecular cell. 2006;23(2):231–239.

76. Krohn M, Wagner R. Transcriptional pausing of RNA polymerase in the presence of guanosine tetraphosphate depends on the promoter and gene sequence. Journal of Biological Chemistry. 1996;271(39):23884–23894.

77. Aivazashvili V, Bibilashvili R, Vartikian R, Kutateladze T. Effect of the primary structure of RNA on the pulse character of RNA elongation in vitro by Escherichia coli RNA polymerase: a model. Molekuliarnaia biologiia. 1981;15(4):915–929.

78. Foster JE, Holmes SF, Erie DA. Allosteric binding of nucleoside triphosphates to RNA polymerase regulates transcription elongation. Cell. 2001;106(2):243–252.

