## Supplementary material for "Approximate Bayesian computation of transcriptional pausing mechanisms": S1 Appendix

### S1 Appendix: Prior distribution justifications

In this appendix we provide brief justifications for the prior distributions selected (Table 1).

Structural studies of the transcription bubble reveal a 10 bp hybrid ($h$) with 0 unpaired nucleotides upstream ($\beta_1$) and downstream ($\beta_2$) [1, 2, 3, 4]. $h$ has also been estimated as 8-9 bp, and the total size of the greater transcription bubble has been estimated as 12-14 bp [5, 6]. We estimated all three of these parameters and conservatively centered our prior distributions around these intervals.

$\lambda_b$ is held constant at 8 nt based off Palangat et al. 1998's [7] estimate of $\lambda_b + h \approx 18$ nt.

The Gibbs barrier of translocation $\Delta G_{\tau}^{\ddagger}$ has previously been estimated as 5 $k_BT$ under the $H_IG_I$ translocation transition model [8]. As we have since normalised the estimates of $\Delta G_{T(l,t)}^{(bp)}$ to control for the transition model, $\Delta G_{\tau}^{\ddagger}$ is recomputed as: $\Delta G_{\tau}^{\ddagger} - 15.96 = 5 \Rightarrow \Delta G_{\tau}^{\ddagger} = 20.96$ $k_BT$. Therefore we centered this parameter around a mean of 21 $k_BT$, and with a standard deviation of 1.

Backtracking and hypertranslocation are slower than translocation between the pre and posttranslocated positions [9, 10]. Our priors for these two processes ($\Delta G_{\tau-}^{\ddagger}$ and $\Delta G_{\tau+}^{\ddagger}$) were set conservatively: with a mean of 5 $k_BT$ and a standard deviation of 2 $k_BT$.

Previous estimates for $k_U$ (0.23 $s^{-1}$ [5]), $k_A$ (3 $s^{-1}$ [5]), and $k_{cleave}$ (0.007 [11] and 0.012 $s^{-1}$ [10]) are sparse. For simplicity, we used the same lognormal prior distribution for all three parameters centered around this interval (with a central 95% interval of (0.007, 3)).

$\lambda_{cleave}$ was held constant at 10 nt based off previous use of the term [10].

In vivo transcription elongation rates of RNAP range considerably, but have been seen as high as 118 bp/s [12]. In practice, $k_{cat}$ is usually estimated to be much greater than the mean velocity [8, 13]. Therefore we set $k_{cat} = 200$ $s^{-1}$. $k_{bind}$ and $\frac{k_{rel}}{k_{bind}}$ were set roughly based off previous estimates for a kinetic NTP binding model [8]. The four NTP concentrations were held constant at *in vivo* estimates made by Traut 1994 [14].

### References


[1] Guo X, Myasnikov AG, Chen J, Crucifix C, Papai G, Takacs M, et al. Structural basis for NusA stabilized transcriptional pausing. Molecular cell. 2018;69(5):816–827.

[2] Kang JY, Olinares PDB, Chen J, Campbell EA, Mustaev A, Chait BT, et al. Structural basis of transcription arrest by coliphage HK022 Nun in an Escherichia coli RNA polymerase elongation complex. Elife. 2017;6:e25478.

[3] Vassylyev DG, Vassylyeva MN, Perederina A, Tahirov TH, Artsimovitch I. Structural basis for transcription elongation by bacterial RNA polymerase. Nature. 2007;448(7150):157.

[4] Vassylyev DG, Sekine Si, Laptenko O, Lee J, Vassylyeva MN, Borukhov S, et al. Crystal structure of a bacterial RNA polymerase holoenzyme at 2.6 Å resolution. Nature. 2002;417(6890):712.

[5] Maoiléidigh DÓ, Tadigotla VR, Nudler E, Ruckenstein AE. A unified model of transcription elongation: what have we learned from single-molecule experiments? Biophysical journal. 2011;100(5):1157–1166.

[6] Nudler E, Mustaev A, Goldfarb A, Lukhtanov E. The RNA–DNA hybrid maintains the register of transcription by preventing backtracking of RNA polymerase. Cell. 1997;89(1):33–41.

[7] Palangat M, Meier TI, Keene RG, Landick R. Transcriptional pausing at+ 62 of the HIV-1 nascent RNA modulates formation of the TAR RNA structure. Molecular cell. 1998;1(7):1033–1042.

[8] Douglas J, Kingston R, Drummond A. Bayesian inference and comparison of stochastic transcription elongation models. bioRxiv. 2018; p. 499277.

[9] Bai L, Shundrovsky A, Wang MD. Sequence-dependent kinetic model for transcription elongation by RNA polymerase. Journal of molecular biology. 2004;344(2):335–349.

[10] Lisica A, Engel C, Jahnel M, Roldán É, Galburt EA, Cramer P, et al. Mechanisms of backtrack recovery by RNA polymerases I and II. Proceedings of the National Academy of Sciences. 2016; p. 201517011.

[11] Saba J, Chua X, Mishanina TV, Nayak D, Windgassen TA, Mooney RA, et al. The elemental mechanism of transcriptional pausing. bioRxiv. 2018; p. 422220.

[12] Ryals J, Little R, Bremer H. Temperature dependence of RNA synthesis parameters in Escherichia coli. Journal of bacteriology. 1982;151(2):879–887.

[13] Abbondanzieri EA, Greenleaf WJ, Shaevitz JW, Landick R, Block SM. Direct observation of base-pair stepping by RNA polymerase. Nature. 2005;438(7067):460–465.

[14] Traut TW. Physiological concentrations of purines and pyrimidines. Molecular and cellular biochemistry. 1994;140(1):1–22.
