## Supplementary material for "Approximate Bayesian computation of transcriptional pausing mechanisms": S2 Appendix

### S2 Appendix: Approximate Bayesian Computation

We used the rejection approximate Bayesian computation algorithm (R-ABC) [1, 2] to infer parameters and select models for the kinetic model classifier. This is in contrast to the more complex Markov chain Monte Carlo approximate Bayesian computation algorithm (MCMC-ABC) we used previously [3]. R-ABC is simple, easy to parallelise, and works as follows:

1. Sample the parameters and model $\Theta_i$ from the prior distribution.
2. Simulate transcription on each gene using $\Theta_i$.
3. Compute the AUC, $AUC_i$, by comparing the simulated pause times with the known positions of pause sites.
4. Accept $\Theta_i$ into the posterior distribution if $1 - AUC_i \leq \epsilon$.
5. Repeat steps 1-4 many times.

Threshold $\epsilon$ is selected to be sufficiently small to ensure that the posterior distribution is an accurate approximation of the true posterior distribution (hence *approximate* Bayesian computation). However $\epsilon$ must also be great enough to obtain an adequately sized sample of the posterior distribution within computational limitations.

Our selection of $\epsilon = 0.255$ is such that the set of models in the posterior distribution has significantly decreased from an entropy of $\log_2(128) = 7$ bits (S2 Fig. 1), is only slightly less adequate than the minimum 1-AUC obtained in the simulations (0.243), and has an adequate sample size ($n = 409$). Our overfitting analysis (Table 2) suggests that increasing $\epsilon$ would not further refine the posterior distribution without increasing overfitting.

The parameter estimates presented in Fig 4 are conditional on the set of models which use the parameter. $\Delta G^{\ddagger}_{\tau+}$ is conditional on $\mathbb{HT} = 1$, $k_U$ and $k_A$ are conditional on $\mathbb{IS} = 1$, $\Delta G^{\ddagger}_{\tau-}$ and $k_{cleave}$ are conditional on $\mathbb{BT} = 1 \vee \mathbb{GT} = 1$, and the other parameters are not conditional on any models.

Parameter estimates are summarised with 95% highest posterior density (HPD) intervals and the geometric median. HPD intervals were calculated conditional on the set of models which use the respective parameter (as indicated above) using Tracer [4]. The geometric median is defined as the posterior sample which has the minimum average Euclidean (for continuous parameters) and Hamming (for discrete parameters) distance from all other posterior samples. Continuous parameters are normalised into z-scores first. The geometric median is calculated conditional on the set of models which use all of the parameters: $\mathbb{HT} = 1 \wedge \mathbb{IS} = 1 \wedge (\mathbb{BT} = 1 \vee \mathbb{GT} = 1)$.

For computational efficiency, the model and parameter estimates presented in Fig 3 and Fig 4 (with the exception of $h$ and $\beta_2$) were partially sampled conditional on the estimates of $h$ and $\beta_2$ derived from the full posterior distribution. $n = 424$ posterior states were used to establish estimates $\hat{h} = 11$ nt and $\hat{\beta_2} = 0$ nt, where $P(h = 11, \beta_2 = 0|D) = 1.0$, and an additional $n = 862$ samples were conditional on these estimates.

The sequence logos in Fig 5C and D were generated by simulating transcription on the test set (100 times for each sequence) using the kinetic model specified by the geometric median and the median time at each transcript length was recorded. The threshold $\theta$ used to classify pause sites was set such that only 1 % of positions are classified as pauses. The sequence logos were then generated from the true positive and false negative pause sites, respectively.

Our code is open source and available at http://www.polymerase.nz.

### References


[1] Beaumont MA. Approximate Bayesian computation in evolution and ecology. Annual review of ecology, evolution, and systematics. 2010;41:379–406.

[2] Csilléry K, Blum MG, Gaggiotti OE, François O. Approximate Bayesian computation (ABC) in practice. Trends in ecology & evolution. 2010;25(7):410–418.

[3] Douglas J, Kingston R, Drummond A. Bayesian inference and comparison of stochastic transcription elongation models. bioRxiv. 2018; p. 499277.

[4] Rambaut A, Drummond A. Tracer 1.6. University of Edinburgh, Edinburgh. UK. Technical report; 2013.

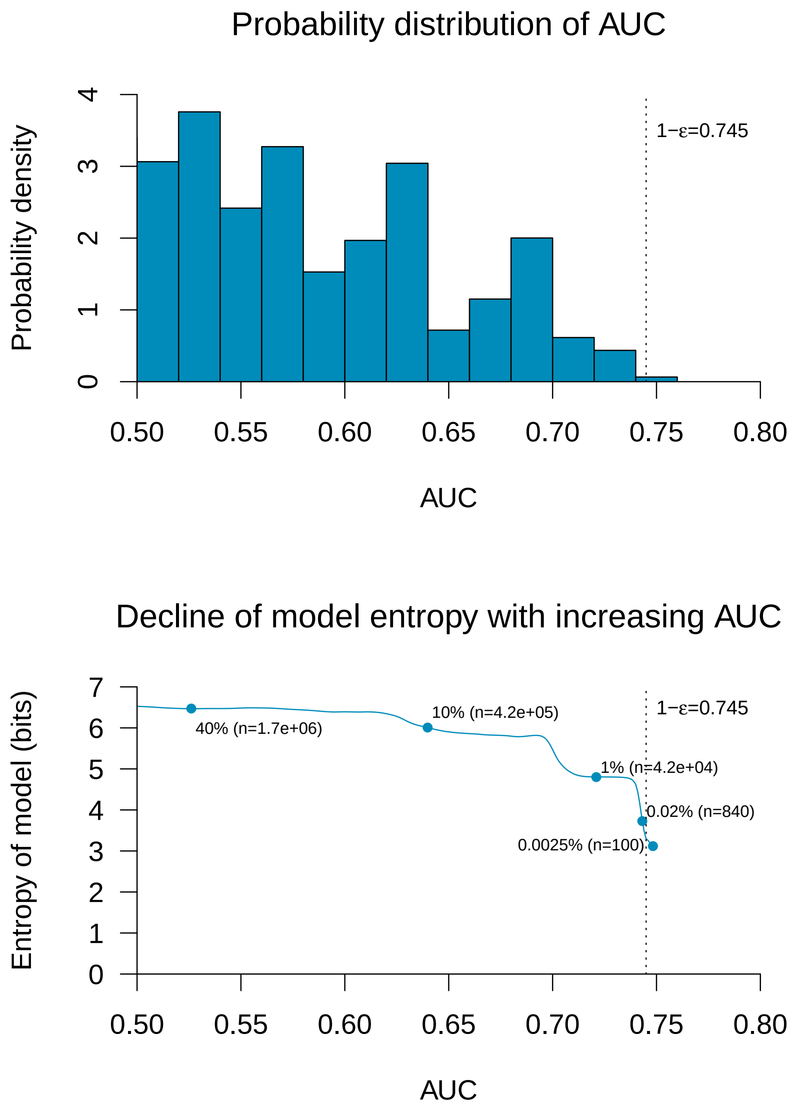


S2 Fig. 1: **Selecting the threshold $\epsilon$.** Top: Probability distribution of $\epsilon = 1-$AUC. Bottom: By decreasing $\epsilon$ (and therefore increasing the minimum required AUC), the entropy of the model features decrease from 7 bits. ABC threshold $\epsilon$ is associated with a significant decrease in entropy, and further decreasing the entropy would only offer minor improvements to the AUC at the cost of $\approx 10^1 - 10^4$ times the computational resources.
